# A computational model of direction selectivity in Macaque V1 cortex based on dynamic differences between ON and OFF pathways

**DOI:** 10.1101/2021.10.23.465582

**Authors:** Logan Chariker, Robert Shapley, Michael Hawken, Lai-Sang Young

**Affiliations:** School of Natural Sciences, Institute for Advanced Study, Princeton, NJ 08540; Center for Neural Science, New York University, New York, New York 10003; Courant Institute of Mathematical Sciences, New York University, New York, New York 10012; School of Mathematics, Institute for Advanced Study, Princeton, NJ 08540

**Keywords:** motion perception, direction selectivity, mechanisms, primary visual cortex, ON-OFF pathways, computational model

## Abstract

This paper is about neural mechanisms of direction selectivity (DS) in Macaque primary visual cortex, V1. DS arises in V1 layer 4Cα which receives afferent input from the Magnocellular division of the Lateral Geniculate Nucleus (LGN). LGN itself, however, is not direction-selective. To understand the mechanisms of DS, we built a new computational model (DSV1) of 4Cα. DSV1 is a realistic, large-scale mechanistic model that simulates many V1 properties: orientation selectivity, spatial and temporal tuning, contrast response, and DS. In the model, DS is initiated by the dynamic difference of OFF and ON Magnocellular cell activity that excites the model’s layer 4Cα the recurrent network has no intra-cortical direction-specific connections. In experiments – and in DSV1 -- most 4Cα Simple cells were highly direction-selective but few 4Cα Complex cells had high DS. Furthermore, the preferred directions of the model’s direction-selective Simple cells were invariant with spatial and temporal frequency, in this way emulating the experimental data. The distribution of DS across the model’s population of cells was very close to that found in experiments. Analyzing DSV1, we found that the dynamic interaction of feedforward and intra-cortical synaptic currents led to cortical enhancement of DS for a majority of cells. In view of the strong quantitative agreement between DS in data and in model simulations, the neural mechanisms of DS in DSV1 may be indicative of those in the real visual cortex.

**Significance Statement:** Motion perception is a vital part of our visual experience of the world. In monkeys, whose vision resembles that of humans, the neural computation of the direction of a moving target starts in the primary visual cortex, V1, in layer 4Cα that receives input from the eye through the Lateral Geniculate Nucleus (LGN). How Direction-Selectivity (DS) is generated in layer 4Cα is an outstanding unsolved problem in theoretical neuroscience. In this paper, we offer a solution based on plausible biological mechanisms: We present a new large-scale circuit model in which DS originates from slightly different LGN ON/OFF response time-courses and is enhanced in cortex without the need for direction-specific intra-cortical connections. The model’s DS is in quantitative agreement with experiments.

## Introduction

This paper is about Direction Selectivity (DS) in the primary visual cortex, V1, of Macaque monkeys. DS has been found in the first stage of V1 signal processing, in neurons in the input layer 4Cα [Hawken and Parker 1984; Hawken et al., 1988; Saul et al 2005]. However, the Magnocellular cells of the Lateral Geniculate Nucleus (LGN) that provide visual input to layer 4Cα are not direction-selective [Wiesel and Hubel 1966; Kaplan and Shapley 1982; Hicks et al 1983; Derrington and Lennie 1984], implying that DS is an emergent property in the cortex.

Understanding the neurobiological mechanisms that generate DS requires a theory to account for the initiation of DS, and, equally if not more challenging, implementation of the theory in a realistic large-scale dynamic circuit model of layer 4Cα that captures the functional properties of the different subpopulations of excitatory and inhibitory cortical neurons. It is well known that DS can result when signals from different locations in the visual receptive field of a neuron have different time-courses [Reichardt 1961], a property known as spatio-temporal inseparability (STI) [Watson and Ahumada 1985; Adelson and Bergen 1985]. There have been many theories of how STI could be produced in cortex, but most have relied on biological mechanisms that are not plausible for Macaque V1. A new theory, based on the different timing of OFF/ON LGN inputs, was proposed in Chariker et al [2021]. Here we implement it in a realistic model (DSV1) of the V1 cortical network to determine whether or not the model can replicate quantitative aspects of DS in V1 neurons.

DSV1 is a large-scale model of Layer 4Cα and its inputs. It is a recurrent, excitatory-inhibitory model in which the choice of most model parameters is guided by experimental data. Like its predecessors [Chariker et al 2016, 2020], DSV1 simulates many of the basic visual response properties of Macaque V1: firing rates, orientation tuning, spatial frequency tuning, contrast response. Here we examine the model’s DS and compare it with that of experimental data obtained in Layer 4Cα.

An innovation in DSV1 is the ON/OFF hypothesis of Chariker et al [2021], namely that ON and OFF Magnocellular LGN cells have small differences in response time-course that initiate DS in the feedforward input to Layer 4Cα cells. A strong point in favor of the hypothesis is that DS is found at the earliest stage of visual processing, in Simple cells in Layer 4Cα that are known to receive Magnocellular LGN input [Chatterjee and Callaway 2003]. Second, the ON/OFF hypothesis achieves STI in Simple cells by using the well-known fact that ON and OFF LGN cells project to distinct sub-regions of Simple cell receptive fields [Hubel and Wiesel 1962; Reid and Alonso 1995]. It is not obvious, however, that a model that incorporates the ON/OFF hypothesis will be able to account for the DS observed in V1 neurons. An influential view of cortical circuit function is predicated on the view that feedforward input is insufficient to generate significant firing rates of V1 neurons because LGN input current is a small fraction of intra-cortical currents in V1 [Douglas and Martin 2004]. Therefore, we had to address the following two crucial questions:

a. are the conditions required for the initiation of DS in [Chariker et al 2021] implementable in a biologically-realistic, recurrent V1 model?
b. is the DS in feedforward LGN inputs alone sufficient for initiating enough DS in the DSV1 model to account for the DS seen across the population of real 4Cα neurons?

The questions are answered at the beginning of the **Discussion**.

## Materials and Methods

### Experimental Methods

#### Animal Preparation

Adult male old-world monkeys (M. fascicularis) were used in acute experiments in compliance with National Institutes of Health and New York University Animal Use Committee regulations. The animal preparation and recording were performed as described in detail previously [Ringach et al. 2002; Henry et al., 2013]. Anesthesia was initially induced using ketamine (5-20 mg/kg, i.m.) and was maintained with isofluorane (1-3%) during venous cannulation and intubation. For the remainder of the surgery and recording, anesthesia was maintained with sufentanil citrate (6-18 μg/kg/h, i.v.). After surgery was completed, muscle paralysis was induced and maintained with vecuronium bromide (Norcuron, 0.1 mg/kg/h, i.v.) and anesthetic state was assessed by continuously monitoring the animals’ heart rate, EKG, blood pressure, expired CO_2_, and EEG.

After the completion of each electrode penetration, 3–5 small electrolytic lesions (3 μA for 3 s) were made at separate locations along the electrode track. At the end of the experiments, the animals were deeply anesthetized with sodium pentobarbital (60 mg/kg, i.v.) and transcardially exsanguinated with heparinized lactated Ringer’s solution, followed by 4 L of chilled fresh 4% paraformaldehyde in 0.1 M phosphate buffer, pH 7.4. The electrolytic lesions were located in the fixed tissue and electrode tracks were reconstructed to assign the recorded neurons to cortical layers as described previously [Hawken et al., 1988]. In 49 animals, more than 700 V1 neurons in all cell layers were recorded in oblique penetrations. For this paper, a full data set was obtained for 94 neurons localized to layer 4Cα.

#### Characterization of visual properties of V1 Neurons

We recorded action potentials (spikes) extracellularly from single units in V1 using glass-coated tungsten microelectrodes. Each single neuron was stimulated monocularly through the dominant eye (the non-dominant eye occluded). Receptive fields were located in the visual field between 1 and 6^0^ from the center of gaze. Stimuli were displayed at a screen resolution of 1024×768 pixels, a refresh rate of 100 Hz, and a viewing distance of 115 cm on either a Sony Trinitron GDM-F520 CRT monitor (mean luminance 90-100 cd/m^2^) or an Iiyama HM204DT-A CRT monitor (mean luminance 60 cd/m^2^). Monitor luminance was calibrated using a Photo Research PR-650 spectroradiometer and linearized via a lookup table.

The response to drifting gratings was used to characterize visual response properties. In this paper, we report measurements of orientation tuning, spatial frequency (SF) and temporal frequency (TF) tuning. We used the f0 response for neurons (traditionally called Complex cells) with f1/f0 ratios < 1 and the f1 response for neurons (traditionally called Simple cells) with f1/f0 ratios > 1. In the population statistics, we counted, for each TF and SF, only cells for which visually driven response exceeded 4 spikes/s, a condition satisfied by most of the cells recorded. All fitting was done using the Matlab function fmincon (Matlab) where the least-squared error was used to minimize the objective function.

#### Orientation tuning

The responses of each neuron were recorded to different orientations between 0 – 360^0^, either in 20 or 15^0^ steps. The stimuli were achromatic gratings at the preferred SF and TF, at a contrast of 64% or greater. All stimuli were presented in a circular window confined to the classical receptive field (CRF).

#### SF tuning

Each neuron was presented with a range of spatial frequencies, usually in ½ octave steps from 0.1 c/deg to around 10 c/deg. For some neurons, the upper limit was extended if the neurons had responses to higher spatial frequencies. We measured SF tuning at the preferred orientation and drift-direction as well as at the non-preferred drift direction.

#### TF tuning

Each neuron was presented with a range of temporal frequencies, usually in 1 octave steps from 0.5 Hz to 32 Hz. Measurements were made for drifting gratings at the optimal orientation in the preferred and non-preferred directions.

## DSV1 Model

The main components of the DSV1 model are shown in Figure 3A. A visual stimulus is provided as input to the model, and we represent it by a light intensity map *L(x,t)* where *x* is a location in 2D visual space and *t* is a time. Magnocellular ON/OFF LGN cells, modeled as leaky integrate and fire neurons, receive input current equal to a spatiotemporal filter of L(x,t), and spikes generated by the LGN neurons then provide feedforward input to the neurons in layer 4Cα. The 4Cα component of the model consists of a network of excitatory and inhibitory leaky integrate and fire neurons arranged in a 2D lattice spanning 1.5 × 1.5mm^2^ of cortex. The corresponding region of L6 of V1, known to feedback to L4 (Callaway, 1998), is also included in the model. Signal transmission from LGN to L4 is assumed to be feedforward only, whereas the interaction between L4 and L6 is bidirectional. We take the part of V1 and LGN modeled to be from ∼ 5 deg eccentricity from fovea, consistent with the cells to be compared within our experimental data. More details on the individual components of the model are given below.

### Visual stimuli and LGN dynamics

#### Visual stimuli

In this paper the visual stimuli were drifting sinewave gratings. The light intensity map *L*(*x,t*) for a drifting grating has the form

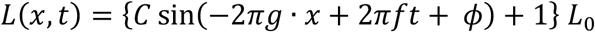

where *x* = (*x*_1_, *x*_*2*_) is a point in the 2D visual space, *t* is time (in sec), *g* is spatial frequency (in c/d) in the direction of the grating, *f* is temporal frequency (in Hz), *ϕ* is the initial phase (in radians), *L*_0_ is the mean light intensity and *C* is the contrast. In the model simulations in this paper, C was set to high contrast as in the experiments.

#### Inputs to LGN cells

The input current to an LGN cell the center of whose receptive field is located at *x*_*0*_ is given by

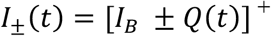

where

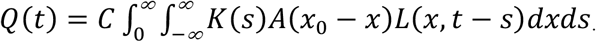

Here the ±sign is for ON and OFF-cells respectively, *I*_*B*_ is background current, *K*(*s*)and *A*(*x*) are the temporal and spatial kernels of the LGN cell, *C* is a constant to be adjusted, and [ ]^+^ denotes the maximum of the bracketed value and 0.

#### LGN Spatial kernel

The spatial kernel is a difference of Gaussians

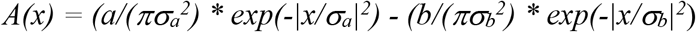

with a=1.0, b=0.74, σ_*a*_=0.0894, σ_*b*_=0.1259 (Zhu et al 2009); *σ*_*a*_ and *σ*_*b*_ are chosen to make the center part of the receptive field correspond to a Gaussian with std. dev. ∼ 0.05^0^ (Derrington and Lennie, 1984). The preferred spatial frequency of the spatial kernel *A(x)* is 2.5 c/deg.

#### LGN Temporal kernels

Each LGN cell is assigned a temporal kernel *K(t)*. All OFF-cell kernels are identical and have the form (adapted from Zhu et al, 2009)

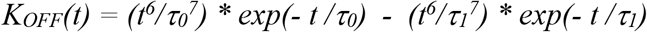

with τ_0_ = 3.66 ms and τ_1_ = 7.16 ms. Parameters are chosen following guidance from temporal kernels derived in experiments (Reid and Shapley, 2002) and the temporal frequency response of LGN cells, ensuring the peak response occurs at ∼10 Hz (Derrington and Lennie, 1984). Note that *K*_*OFF*_ is positive before the zero crossing, as the polarity of the LGN cell is implemented in the sign of +/- in the expression for I(t), not in the kernel.

Differences between ON and OFF temporal kernels are introduced (following Chariker et al, 2021) in order to produce directional preference in the input received from LGN cells (see **Results**). A variety of ON cell temporal kernels is used in the model following the diversity in ON cell temporal kernels sampled experimentally (Reid and Shapley, 2002). Using the notation (a,b) to denote the temporal kernel *K*_*a,b*_ defined in **Results**, the table below shows the distribution of kernels given to the ON cells.

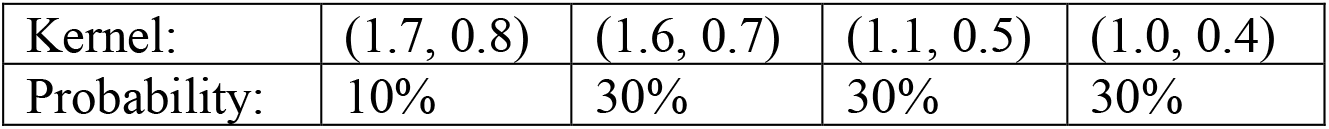

A (1,1) kernel is identical to the OFF cell kernel *K*_*OFF*_, and an (a,b) kernel is equal to *K*_*OFF*_ multiplied by *a* before the zero crossing and by *b* after the zero crossing. Each of the above kernels has a/b approximately equal to 2, which is shown in Chariker et al (2021) to produce direction selectivity in the feedforward input to V1 at temporal frequencies below ∼ 6 Hz. ON cell kernels are also given a delay of 9-11 ms, shown to produce DS in the feedforward input to V1 at temporal frequencies above ∼ 4 Hz (see **Results** and Chariker et al, 2021).

For ½ of the model ON cells, the part of the temporal kernel after the zero crossing is stretched out in time to reduce the total kernel area to 0, as many experimentally sampled ON cell temporal kernels are approximately biphasic (Reid and Shapley, 2002). We found that at low temporal frequencies (4 Hz and below) adjusting the area to zero in this way significantly reduces the direction selectivity in the feedforward input to V1 cells, and consequently we left ½ of the ON cell kernels unchanged. Further details are given in **Supplementary Information** (S1).

#### Dynamic equations

LGN responses are modeled with noisy-leaky-integrate-and-fire (NLIF) equations with *I*±(*t*)as input [Lin et al 2012]. The membrane potential *V* of an LGN neuron is assumed to satisfy

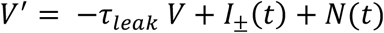

with spiking threshold at ∼1 and reset to 0. Here τ_*leak*_ is a constant and *N(t)*is a noise term. As the firing rates of LGN cells are known to saturate under strong drive (Kaplan et. al, 1997), we limit the ability of a model LGN cell to produce spikes in rapid succession by adding a brief increase to the spiking threshold immediately after a spike. This has the effect of lowering LGN firing rate when strongly driven more than when weakly driven. Implementing this mechanism causes the LGN firing rate response as a function of temporal frequency to more closely match experimental data (Levitt et. al., 2001). Further details are given in **Supplementary Information** (S1).

### Cortical components

#### Cortical connections in DSV1

The primary focus of the DSV1 model is on the feedforward inputs from LGN --> Layer 4Cα (L4), the input layer to V1 in the Magnocellular stream, and the interactions of these inputs with intra-cortical dynamics. A third component of DSV1 is Layer 6 (L6) of V1, with which L4 is known to interact [Callaway 1998]. DSV1 is descended from models in [Chariker et al 2016, 2018, 2020]: mechanistic, computational models of cortical function and mechanisms. The model in [Chariker et al 2020] replicates successfully many functions of V1, such as orientation and spatial frequency selectivity, contrast response, the production of gamma rhythms, and the presence of Simple and Complex cells. Its neurons do not, however, have direction selectivity. DSV1 deviates substantially from its predecessors in having different dynamics of the modeled LGN ON and OFF cells, and in how the LGN cells are connected to V1 neurons, as this is where DS in the model originates. This part of the model is discussed in detail in **Results**. The modeling of L4 and L6 in DSV1 is quite similar but not identical to [Chariker et al 2020]. We first recall the main features of DSV1 that are largely unchanged before discussing what is different.

#### Dynamics of 4Cα neurons

Layer 4C*α* consists of a network of excitatory (E) and inhibitory (I) neurons with membrane potentials evolving according to a set of leaky integrate and fire (LIF) equations. Potentials are denoted by 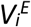 and 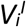 for E and I neurons, respectively, with *i*=neuron index. Neurons spike when *V* reaches a threshold potential of *V*_*thresh*_*∼1*. Upon reaching the threshold, *V* is instantaneously reset to a resting potential value of *V*_*rest*_ = 0, remaining for an absolute refractory period of *t*^*E*^_*ref*_ = 2 ms for E and *t*^*I*^_*ref*_ = 1 ms for I. While not in the refractory period, *V* obeys the Leaky-Integrate-and-Fire (LIF) equation [Knight 1972]:

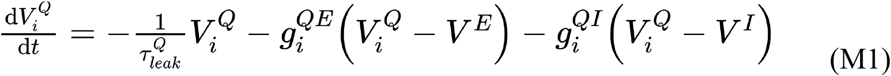

where Q=E or I denotes the neuron type; V^E^=14/3 and V^I^=-2/3 are the excitatory and inhibitory reversal potentials; and *g*^*QE*^ and *g*^*QI*^ are the excitatory and inhibitory conductances. *g*^*QE*^ is a sum over all excitatory postsynaptic conductances (EPSC) generated by the presynaptic excitatory spikes of layer 4Cα, layer 6, and LGN, as well as from independent, Poisson-timed kicks modeling the effects of ambient neurotransmitter. *g*^*QI*^ is a sum of inhibitory postsynaptic conductances (IPSC) from the presynaptic inhibitory spikes in layer 4Cα. In the case of Q=E, an additional slow time course IPSC is generated from each presynaptic inhibitory spike. Details on the EPSC and IPSC time courses can be found in Chariker et al [2018].

#### Layer 4Cα (L4)

We modeled 9 hypercolumns (HC) of 4Cα. Each HC measures 0.5 × 0.5 mm and is divided into regions within which cells are intended to have similar orientation preferences. Following [Beaulieu et al 1992], we used cell densities of ∼4000 neurons per HC, 3/4 of which are E cells and the rest are I cells. The I-neurons are assumed to be a homogeneous population of local-circuit basket cells, a reasonable approximation for layer 4Cα [DeFelipe et al 1999]. E-and I-neurons are uniformly distributed in the model cortex, and their dynamics are governed by conductance based integrate-and-fire equations. The probability of connection between model cells is dependent on distance in the cortex and on cell types (E or I), while the strength of connection is independent of distance [based on Oswald and Reyes 2011]. For presynaptic E-neurons, the distance-dependence of the connection probabilities are given by Gaussians with standard deviation 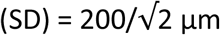; for presynaptic I-cells, 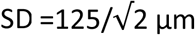 [Fitzpatrick et al 1985; Yoshioka et al 1994]. Peak connection probability between E-cells is ∼15% on average. I-cell connections are denser. Following Holmgren et al [2003] and Oswald and Reyes [2011], we set peak I-to-I connection probability at cortical distance=0 to be 60%.

The numbers of connections and cell densities imply that on average, an E-cell has ∼200 presynaptic E-cells and ∼100 presynaptic I-cells; for an I-cell, the corresponding numbers are ∼750 E-cells and ∼100 I-cells. In addition to cortical inputs, layer 4Cα E- and I-cells receive monosynaptic input from 1-6 Magnocellular LGN neurons (more details given in **Results**).

#### Feedback from Layer 6 (L6)

In DSV1 as in the real cortex, L4 and L6 interact continuously in a self-adjusted feedback loop. The details of the L4-L6 feedback loop were specified in Chariker et al [2020]. The goal of Chariker et al [2020] was to construct a model that could simulate the response of V1 over a wide range of stimulus contrast, and the L4-L6 feedback loop played a large role in model performance. In this paper the model was used to simulate only experiments that employed high contrast stimuli and thus the L4-L6 feedback loop was working only at one contrast level throughout. We refer the reader to Chariker et al [2020] for a complete documentation of the model details given in the SI of that paper.

### Two new features of DSV1

#### 1) Firing rate homeostasis

Without intervention, firing rates of Simple E-cells in DSV1 varied widely. Homeostatic mechanisms that could support adjustment of synaptic input to stabilize firing rates are known to be present in cortex [Turrigiano 2012]. Homeostasis was introduced into DSV1 by giving Simple E-cells in L4 that are very high firing more I-input (e.g. 5-10 more I-cell inputs in addition to the usual complement of ∼100 I cell inputs). This produced a small reduction of firing rates in the Simple cells with the highest firing rates. It also had the effect of reducing an unwanted correlation in DSV1 between high firing rate and low DS, a correlation that was not present in the cortical data.

#### 2) Adaptation at low temporal frequencies

DSV1 is the first in this line of models to consider a full range of TF. To control the build-up of excessive spiking of Complex cells at low TF, it is necessary to incorporate a form of spike rate adaptation. We implemented a scheme to make it harder for an E-neuron to continue to fire after a burst of 7 consecutive spikes are fired within 150 ms. Adaptation mechanisms are also known to be present in the real cortex [Barkai 2005; Adelman et al 2012]. More details are given in **Supplementary Information** (S3).

### Model parameters

The synaptic coupling strengths are an important part of the model. They are set in the same way as in previous models [Chariker et al 2016, 2020] with the use of spike firing data and intracellular recording data as guides. The coupling strength between E cells in L4, termed S_EE_, was set to 0.023, consistent with experimental data [Stratford et al 1996]. The coupling strength from I to E cells, S_EI_, was calculated to be 2.16·S_EE_ based on firing rates and intracellular data [cf. **Materials and Methods** in Chariker et al 2016]. The coupling strength between I cells, S_II_, was set to be 0.65·S_EI_. The remaining coupling strength S_IE_, the coupling strength from E to I, was determined to be approximately 0.26·S_EE_ by simulating a range of values of S_IE_ and finding what value of S_IE_ was compatible with the mean background firing rate of E cells as measured experimentally. Based on the literature, we set the LGN synaptic coupling weights to be S_E,LGN_=2.3·S_EE_ and S_I,LGN_=2.85·S_EE._ The coupling coefficients for L6, based on [Stratford et al 1996] were S_E,L6_=0.36·S_EE_ and S_I,L6_=0.087·S_EE_. See **Supplementary Information** (S5) for details.

### Statistical comparison of the cumulative distribution functions (CDFs) of model and data

We used a bootstrap procedure to calculate the similarity or dissimilarity between the CDF that describes the Pref/Opp ratio of experimental data and the CDF of the cells in the DSV1 model (Results, Fig. 7). We generated 1000 sample distributions of DS (=Pref/Opp) from the model, each having the same number of cells as the data distribution: N=62 in the case of Simple cells and N=32 in the case of complex cells. We used sampling with replacement. Then we calculated the CDF for each sample and the data by binning the sample distributions and the data distribution in the same way. The CDF had 7 bins: 1-1.414, 1.414-2, 2-2.828, 2.828-4, 4-5.66, 5.66-8, and >8. That was done so that each bin of the CDF had O(10) cells. Then we calculated the summed squared deviation (SSD) between each sample CDF and the CDF of the full population of model cells: 2031 Simple cells for the Simple cell CDF and 914 Complex cells for the Complex cell CDF and also the SSDs for the data CDFs for both Simple and Complex cells. The statistical comparison was between the SSDs for the data (Simple and Complex) CDFs and the distribution of SSDs for the 1000 samples for the Simples and 1000 samples for the Complexes. The result was that the SSD for the Simple cell data CDF was 0.91 * St.Dev. for the 1000 samples, and the SSD for the Complex cell data CDF was 0.21 * St.Dev. for the 1000 samples. This leads to p>0.2 for Simple cells and p>0.7 for Complex cells.

## Results

Here is a brief guide to the **Results** section; it contains: (1) data that establish that there is a high proportion of Simple cells in Macaque layer 4Cα that are direction-selective, and the broad-band nature of their DS, (2) the ON/OFF hypothesis of Chariker et al [2021] about DS in the feedforward input to V1, (3) the design of the DSV1 model, (4) a comparison of DS in model and data, and (5) an analysis of the DSV1 model that shows how cortex boosts the DS present in its feedforward inputs.

### Experimental data on DS in Macaque V1, layer 4Cα

Action potentials (spikes) were recorded extracellularly from single units in Macaque V1 using tungsten microelectrodes. We measure responses to drifting, achromatic, sinusoidal gratings to characterize visual response properties (see **Materials and Methods**). Stimulus contrast in these experiments was 64% or greater. In 49 animals, the activity of hundreds of V1 cells in all cell layers was recorded in oblique penetrations; complete data were obtained for ∼700 cells that could be assigned accurately to a cell layer. The data in this paper are from neurons localized to layer 4Cα on the basis of histology (Figure 1A; see **Materials and Methods**). Cortical cells were classified as Simple and Complex cells conventionally, on the basis of the modulation ratios (f1/f0) of their responses to drifting gratings [Skottun et al 1991; Ringach et al 2002; **Materials and Methods**]. We present in Figure 1C-E data for 94 cells in layer 4Cα: 62 Simple cells and 32 Complex cells.

**Figure 1.**
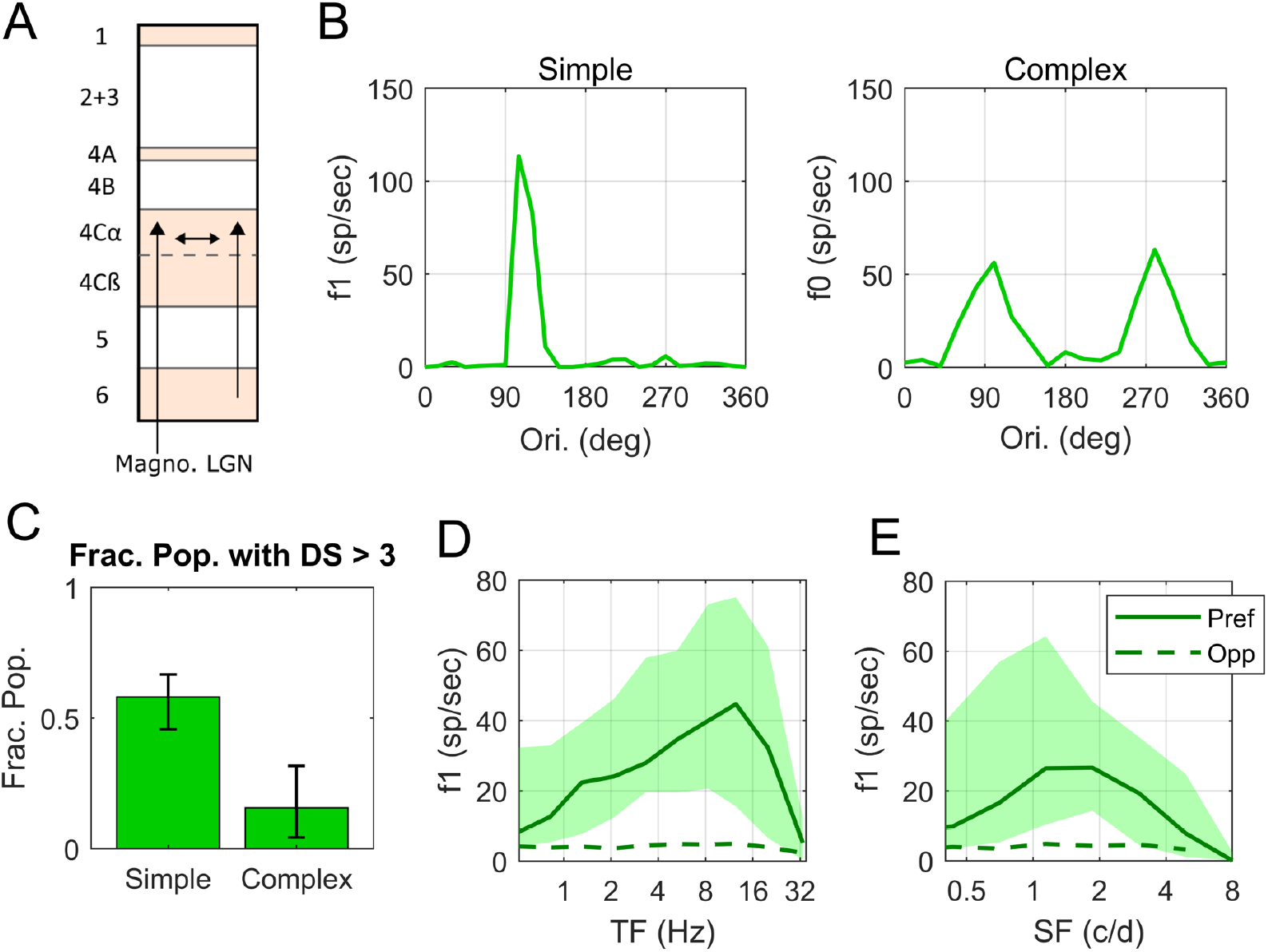
Experimental Data on DS and firing rates as a function of TF in macaque layer 4Cα. **A**. Diagram showing a cross section of the layers of V1, with arrows indicating the recurrent interaction within layer 4Cα, feedforward inputs to layer 4Cα from LGN and feedback to layer 4Cα from layer 6. All data in this figure are from cells recorded in layer 4Cα. **B**. Orientation tuning curves for a highly DS Simple cell and a non-DS Complex cell. **C**. Fraction of Simple and Complex cells with DS (=Pref/Opp) > 3. In our sample, 36 of 62 Simple cells and 5 of 32 Complex cells had DS > 3, with 95% confidence intervals. **D**. Temporal frequency (TF) tuning curves of Pref and Opp firing rates in Simple Cells with DS>3. The thick solid curve shows median Pref firing rate and dashed curve shows median Opp firing rate. Quartiles indicted by the shading around the median. **E**. Spatial frequency (SF) tuning curves; same setup as in **D**. Note that the median rate here is lower because there is a range of preferred spatial frequency across the population therefore the median underestimates the median of the rate at the preferred SF.

A large fraction of the cells recorded in layer 4Cα had a high value of DS. Representative examples of single-cell responses are shown in Figure 1B. The response vs. orientation of one 4Cα Simple cell is shown in the left panel and of one Complex cell in the right panel. The data in the graphs of Fig. 1B are the responses to drifting gratings at the (SF, TF) combination that produces the highest response at the stimulus drift TF (f1) in the Simple cell or the highest mean firing rate (f0) in the Complex cell. The measure of DS that we use throughout the paper for both experimental data and model responses is the ratio Pref/Opp. For Simple cells, Pref is taken to be the f1 response in the orientation of maximal response (for Complex cells we use f0), and Opp for Simple cells is the f1 response at 180° from the Pref direction (for Complex cells again we use f0). Across the population of 4Cα Simple cells in our sample, the fraction with DS>3 is 0.58 (Fig. 1C); the fraction of 4Cα Complex cells with DS>3 is only 0.16 (Fig. 1C) [cf. Hawken et al 1988]. The DSV1 model emulates the much higher incidence of DS in Simple cells, as shown later in **Results**.

An important property of DS in layer 4Cα is the broad-band character of DS (Fig. 1D, E) in Simple cell responses. In the experiments, we measured responses at preferred and opposite-to-preferred directions over a wide range of TF and SF for 53 Simple and 29 Complex cells. The preferred and opposite-to-preferred directions identified at the peak of the ori-tuning curve as in Fig. 1B were the same directions for all TF and SF, i.e the preferred direction remained consistent over the entire range of response. In Fig. 1D, the median Pref (f1) response (solid curve) across the population of 4Cα Simple cells that had high DS (Pref/Opp>3; 26 cells) is much larger than that of Opp (dashed curve) across the full range of temporal frequency (TF) to which the cells respond, not just at or around the peak response near 10 Hz. We also measured DS as a function of SF in the high DS Simple cell population (Fig. 1E; 25 cells). As with TF, Pref responses are larger than Opp over the full range of SF that is effective in driving V1. The broad- band character of DS in 4Cα neurons is reproduced in the DSV1 model, as shown later in **Results**.

### Theory of DS in feedforward inputs and OFF-ON dynamics

First we describe the theory proposed by Chariker et al [2021] that is embedded in the DSV1 model, and then in the following section we describe the model itself. The basic concept developed in [Chariker et al 2021] is that different response time-courses of OFF and ON LGN neurons afferent to a cortical cell would result in DS in their summed responses that comprise the feedforward input to 4Cα neurons.

Reducing two rows of OFF and ON LGN cells to an OFF-ON pair the receptive fields of which are separated by *d* degrees (Figure 2A) and assuming the responses of individual LGN cells are functions of sinusoidal form (a simplification), we analyzed in [Chariker et al 2021] the summed response of the OFF-ON pair to left and right-moving gratings. The key findings can be summarized as follows:

**Figure 2.**
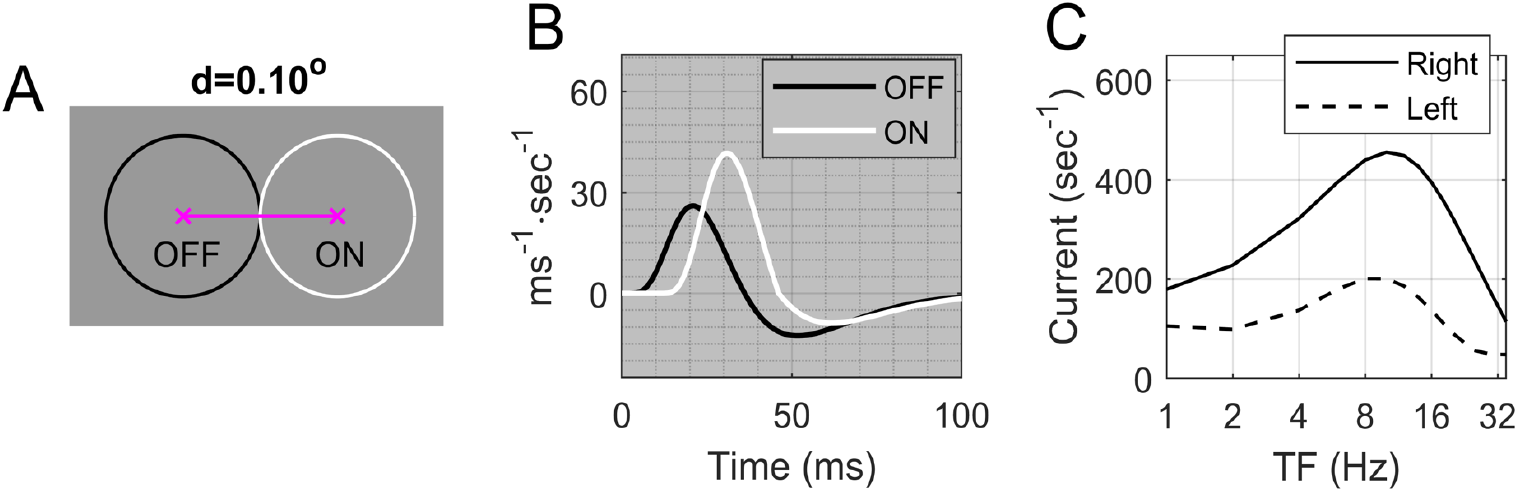
Summed response from a pair of ON and OFF LGN cells. (redrawn from [Chariker et al 2021]). **A.** ON-OFF pair separated by *d*°, with ON to the right of OFF; the circles represent 1 SD of the center Gaussian of each LGN receptive field. **B.** Relative to the OFF-kernel (Black), the ON-kernel (White) has a 10 ms delay and is of type (1.6, 0.7) (see text under Model Description) **C.** Right vs Left responses (summed ON and OFF currents) as functions of TF, at SF=2.5 c/d.

1. DS is equivalent to asymmetric ON/OFF phase differences in response to left and right gratings: the grating that elicits the smaller phase difference (i.e., where ON and OFF elicit firing more in-phase) is the preferred direction. This idea, known from earlier work [Watson and Ahumada, 1983, 1985], is behind much of the theory.
2. Two mechanisms for inducing such asymmetric ON/OFF phase differences are: (i) *delay* in the ON-response and (ii) *different time kernels of ON vs OFF cells*, with the ON-kernel having a taller positive lobe. Mechanism (i) is effective at high TF; mechanism (ii) at low TF. See Fig. 2B. These two mechanisms combined produce a consistently preferred direction, from OFF to ON, with roughly the same amount of DS over a broad range of TF, as illustrated in Fig. 2C.
3. If the ON/OFF separation *d* is smaller than half the period of the stimulus grating, then directional preference of the OFF-ON pair will be consistent over the entire range of SF visible to the LGN pair [Chariker et al 2021, Fig. 5]. This means, for example, that values of *d* should be < 0.10 deg of visual angle for an eccentricity of ∼5°. In general, optimal SF, receptive field size, and *d* scale with eccentricity.

The findings above describe how DS in summed ON-OFF LGN response is produced in an idealized setting. It is understood that there will be much more variability in the real brain.

### DSV1 Model description

The general physical layout of the DSV1 model is shown in Figure 3A. The model has three components: Magnocellular LGN cells, Layer 4Cα (L4) and Layer 6 (L6) of the primary visual cortex (V1) (cf. Fig. 1A). The region of cortex modeled represents an area of roughly 0.75 × 0.75° on the retina located at about 5° eccentricity. Our primary focus is L4, populated by 36,000 Excitatory (E) and Inhibitory (I) neurons, coupled with synapses that have realistic conductances. All the neurons in the model obey the Leaky Integrate-and-Fire (LIF) differential equation [Knight 1972] written in **Materials and Methods**. The DSV1 model, like its predecessors [Chariker et al 2016, 2020] is highly recurrent, consistent with the modern view of visual cortical function [Douglas and Martin 2004]. The model cortex gets all its LGN input from one eye only; the inclusion of ocular dominance columns would not impact the character of the feedforward input from LGN to cortical cells.

**Figure 3.**
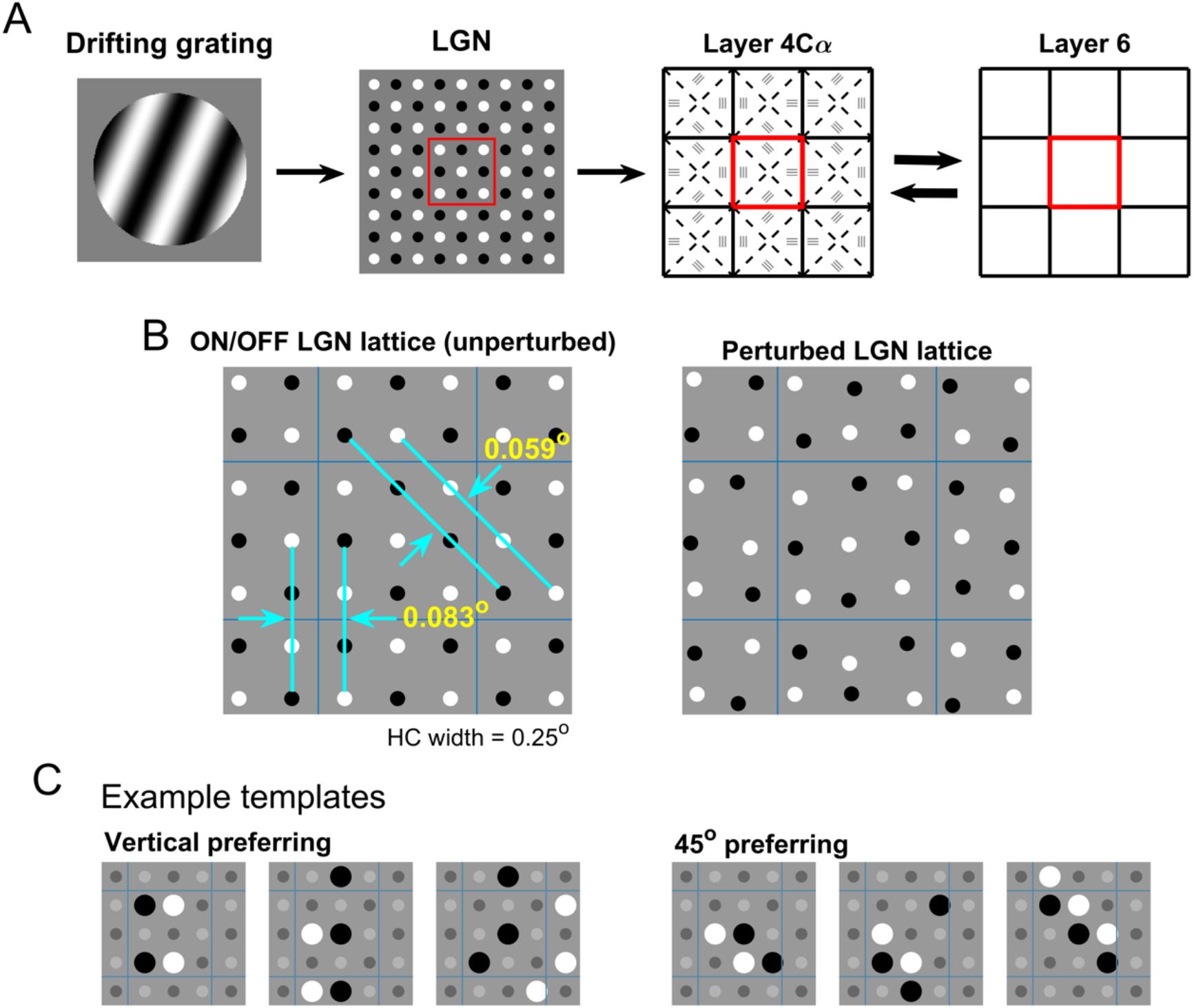
The DSV1 model and its LGN templates. **A**. General layout of the three components of the DSV1 model. The visual stimulus is represented by the drifting grating on the left. This information goes to the sheet of Magnocellular LGN cells. Each cell computes a response R(t) as described in **Results**. LGN outputs are then passed to Layer 4Cα of V1, the primary input layer of the Magno stream. Layer 4Cα receives feedback from Layer 6. **B**. LGN sheet showing ON-OFF separation distances: ON cells in white, OFF cells in black; square lattice unperturbed (left), perturbed (right). **C**. Six example LGN templates, three for vertical, and three for 45°, with NLGN= 4, 5, 6.

Inputs to DSV1 are visual stimuli represented as time-dependent light-intensity maps *L(x,t)* where x denotes the location on the retina (or on the LGN sheet) and *t* denotes time. Once *L(x,t)* is presented to the model, LGN computes a response, which is passed to L4, and dynamic interactions within L4 and between L4 and L6 produce a response. Only drifting gratings are used as stimuli in the current study because most experimental data about DS were obtained with such stimuli, but the DSV1 model’s capabilities are not limited to this class of visual stimuli.

DSV1 deviates from its predecessors [Chariker 2016, 2020] in its OFF-ON LGN dynamics and in the design of LGN “templates”, referring to configurations of LGN cells projecting to a cortical neuron. These modifications, which follow prescriptions from [Chariker et al 2021], are necessary to simulate cortical DS; in our earlier models that had ON and OFF LGN cells with the same response dynamics [Chariker 2016, 2020], model neurons were not direction-selective. The architecture of the network, the equations governing the dynamics, and the parameters used are similar though not identical to those in [Chariker 2020]; they are described in detail in **Materials and Methods**. Below we focus on DSV1’s most significant deviations from its predecessors, namely LGN dynamics and thalamo-cortical convergence.

### LGN Dynamics

In DSV1, ON and OFF cells have different dynamics, i.e. visual response time-courses, following the prescription in Chariker et al [2021].

First, we introduce a delay in the response of ON LGN cells with respect to OFF. Such a delay is consistent with retinal circuitry required for the sign inversion in the ON pathway [Masland 2012] and with direct measurements of LGN visual responses [Reid and Shapley 2002; Jin et al 2011]. In DSV1 the time-delay between ON and OFF varied between 9-11 ms (Fig. 2B).

Following Reid and Shapley [2002], we model the temporal kernel shapes of OFF and ON cells as follows: Let the OFF-kernel *K*_*OFF*_*(t)* = *K(t)* be as defined in **Materials and Methods**. Using *K*^*+*^*(t) = max{0, K(t)}* and *K*^*-*^*(t) = min{0, K(t)}* to denote the positive and negative parts of *K(t)*, we consider the family of temporal Kernels *K*_*a,b*_*(t)* defined by

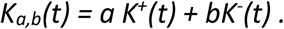

In this notation, *K*_*OFF*_*(t)* has the structure (1,1). It is shown in Chariker et al [2021] that a kernel *K*_*a,b*_*(t)* with *a*>*b* produces a phase lag relative to *K*_*1,1*_*(t)*. In the DSV1 model, ON-cells had temporal kernels with a variety of structures (*a, b*) with *a/b ∼ 2* (consistent with data in [Reid and Shapley 2002]); they range from (1.7, 0.8) to (1.0, 0.4) drawn randomly for each cell. The OFF kernel and a typical ON kernel are illustrated in Fig. 2B.

### Template design

One of the features that distinguishes the DSV1 model and its predecessors [Chariker et al 2016, 2018, 2020] from other existing models of the visual cortex is their realistic depiction of the small number of LGN cells per unit area of the visual field in the M-cell – layer 4Cα pathway. Two independent estimates based on M-retinal ganglion cell density [Silveira and Perry 1991] and on Macaque LGN data [Connolly and Van Essen 1984] point to an average of about 9 LGN cells projecting to an area of 0.25° times 0.25° in the visual field, roughly the retinal projection to one V1 hypercolumn (HC) at 5° eccentricity. Using different inferences, Garcia-Marin et al [2019] suggested a somewhat higher figure for LGN-V1 convergence. Whichever estimate one uses, the M-cell LGN input to V1 cortex is very sparse. Given that the rows of ON and OFF LGN cells cannot be too far apart to ensure a consistent directional preference as seen in data (see Figure 1 and Result (3) in the summary of results from [Chariker et al 2021]), the LGN sparseness poses a significant challenge to modeling.

In DSV1, we model the locations of LGN cells by a perturbed square lattice of alternating ON/OFF cells as shown in Fig 3B. We start from a regular lattice with 9 LGN cells/HC (Fig 3B left), and perturb each cell’s position in the lattice with independent Gaussians with SD = 0.01° resulting in the somewhat irregular mosaic on the right. In the (unperturbed) lattice, there are very few discrete choices of ON-OFF distances. Between two vertical rows, the distances are multiples of 0.083°, and between two diagonal lines making 45° with the vertical, the distances are multiples of 0.059°. Both are within a favorable range of *d* [Chariker et al 2021].

By an LGN “template”, we refer to a configuration of LGN cells that forms the inputs to a cortical cell; translates of the same configuration are regarded as coming from the same template. In the DSV1 model, there are 4 distinct collections of LGN templates; they prefer 0° (which we define to be vertical), 45°, 90° and 135°. The latter two collections are obtainable from the first two by a 90° rotation, so we need only to be concerned with vertical and 45° templates. The number of LGN afferents to a cortical cell in the model is between 1 and 6, with slightly less than half of the cortical cells receiving 5 or 6 LGN inputs. Those numbers are based on the sparseness of LGN together with results from Angelucci and Sainsbury [2006] on thalamo-cortical convergence. The chosen LGN templates consist mostly of configurations where the ON-OFF rows are as close to each other as possible. Several examples of the templates used are shown in Fig 3C. We switched from the perturbed hexagonal lattice used in [Chariker 2016, 2018] because the design of templates with ON/OFF separations < 0.1° is simpler using a square lattice but we duplicated all of the results in this paper using hexagonal lattices (not shown), proving that the results are not tied to the specific lattice choice.

### Connection of LGN cells to V1 cells

Following are the rules for assigning to each V1 cell a group of LGN cells that project to it. In layer 4Cα, we divide each HC of L4 into 4 wedges (Fig. 3A). Cells in each wedge are given the intended orientation indicated by the 3 bars drawn in that wedge, and cells near the boundary of two wedges have probability ½ of picking each of the two intended orientations. There are two main parts to LGN assignment: (a) *Selection of template*. From the collection of templates for that orientation, we picked randomly according to prescribed probabilities (i) the number of LGN cells in the template, and (ii) two vs three ON/OFF subregions. (b) *Admissible locations*. All possible translates of the configuration chosen in (a) that lie within a radius of 0.3° of the V1 cell’s retinal projection are admissible and are chosen with equal probability. Details are given in **Supplementary Information** (S2).

To be clear, we have not built into DSV1 any capability for generating directional preference beyond DS in feedforward LGN inputs: for a cortical cell, presynaptic E and I-cells are isotropically distributed, there are no special connections between cells with the same directional preference, and no long-range intra-cortical lateral connections in layer 4Cα. In local populations of 4Cα cells, LGN templates of opposite parity (in ON/OFF) were drawn with equal probability to avoid introducing biases in preferred directions.

### DS properties of the DSV1 model and comparison with data

#### Time series of membrane voltage and currents in DSV1

Comparing membrane potential and synaptic current traces in the preferred and opposite directions for DS neurons in DSV1 is revealing about the role of LGN inputs in driving cortical dynamics. Figure 4 shows voltage traces for a typical model Simple cell of high DS, in Pref (Fig. 4A) and Opp (Fig. 4B) directions when driven by an optimal stimulus grating of SF=3 c/deg and TF=10Hz. Spike occurrences are indicated in the plots just below the respective voltage plots. Many more spikes are fired in response to the stimulus in the preferred direction and the firing rate in the preferred direction is modulated at the stimulus TF of 10Hz.

**Figure 4.**
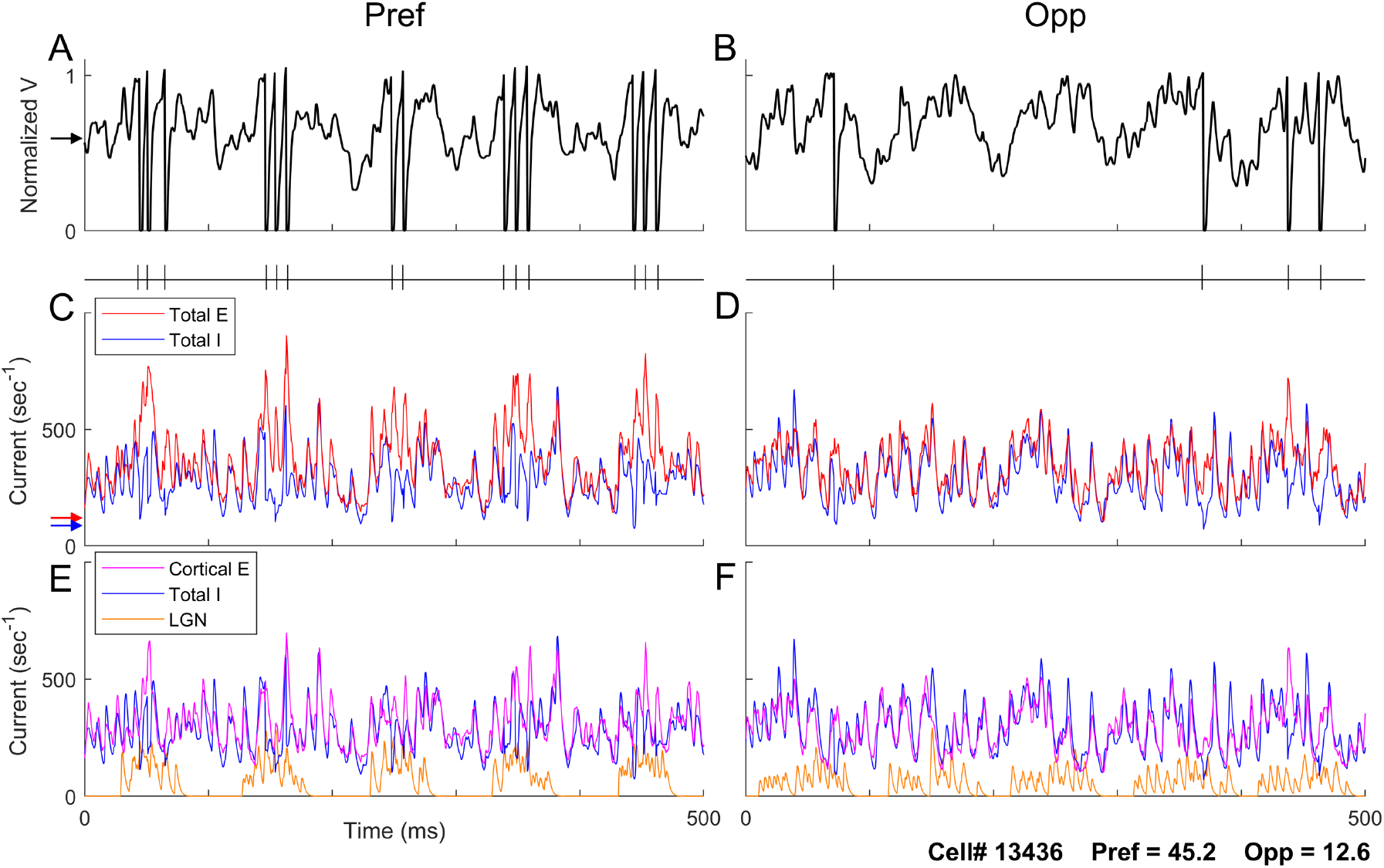
Voltage and input current traces in Pref and Opp directions for a model high DS Simple cell. Responses of a high DS cell in the model to drifting gratings of 3 c/deg and 10 Hz, optimal for this cell. Panels A, C and E are for Pref; B, D and F are corresponding panels for Opp. **A(Pref), B(Opp)** voltage traces and spike times. **C (Pref), D(Opp)**. Total synaptic input currents: Total Excitatory current (= L4+L6+LGN+amb) (red), negative of Inhibitory current (blue). **E(Pref), F(Opp)**. Synaptic input currents by source: LGN current (orange), cortical E-current (= L4+L6+amb) (purple), negative of I-current (blue). Arrows on the left show time averages during *spontaneous activity* of (in **A**) membrane potential (black), (in **C**) Total E (red) and Total I (blue) synaptic currents.

The plots of membrane current illustrate what is happening intracellularly in a model DS cell when it receives visual input. Fig. 4C, D show membrane synaptic currents in Pref and Opp directions. “Total E” here refers to the sum of the E-currents from L4, L6, LGN and ambient sources (see **Materials and Methods** for the definition of “ambient”). Total I is the magnitude of the input I-current, plotted with a positive sign so we can compare the magnitudes of the E and I currents. The increase in current during stimulation can be seen by comparing to the arrows on the left; they show mean E and I-currents during spontaneous activity.

Fig. 4E, F show the decomposition of Total E into LGN and “cortical-E”, referring to L4+L6+amb. The cortical-E and I-currents track each other over time (Fig 4E, F) as was documented in our previous recurrent models [Chariker et al 2018]. The covariation of these currents in gamma-band oscillations is an emergent phenomenon in highly recurrent models like DSV1, producing a moment-by-moment balance that is a stronger form of E-I balance than that in classical balanced state theories [van Vreeswijk and Sompolinsky 1997; Hansel and Sompolinsky 1996]. In the Pref direction (Fig. 4C), the E-I balance is broken roughly every 100 ms (the grating’s TF = 10 Hz), when the Excitatory current is more strongly modulated by its LGN inputs. When that happens, the E-current momentarily overwhelms the I-current causing spikes to be fired. In the Opp direction (Fig. 4D), the LGN drive is smaller and more spread out in time. It is less effective in producing the sudden surges in excess E-current needed to drive the membrane potential over threshold. These current plots show clearly that feedforward LGN inputs in Pref and Opp directions impact cortical firing, even though LGN synaptic current comprises only a small fraction of the total E-current entering a cell.

LGN feedforward inputs have such a large impact on Simple cell spike firing because of the balance of excitatory and inhibitory cortical synaptic currents [Hansel and Sompolinsky 1996]. For a cortical cell in the model, firing rate is roughly determined by the difference between its Total E-current and Total I-current; this is a direct consequence of the Leaky Integrate-and-fire equation (**Materials and Methods**). When cortical-E and I currents are approximately balanced as they are in DSV1 (Fig. 4E, F), each cortical cell’s spike firing is very sensitive to the size of the modulated Feedforward input. The behavior illustrated in Figure 4 is typical for DS cells in the DSV1 model.

### Visual responses of single cells in the data and in the DSV1 model

The tuning curves of individual model cells in DSV1 indicates that their visual properties closely resemble those of typical 4Cα Simple cells in the real cortex (Figure 5). Plotted on the left of Fig. 5 are tuning curves of cells recorded in layer 4Cα (in green); on the right are the tuning curves of model Simple cells (in black). The first row shows orientation tuning, the second row shows TF responses in Pref (solid) and Opp (dashed) directions, and the third row are plots of SF responses. Cells 1 and 2 in the data have high DS. Their orientation tuning curves have only one peak over the full 360^0^ range of orientation, and the Opp responses in the TF and SF tuning curves are much smaller than Pref responses, with no changes in directional preference across TF or SF. Cell 3 in the data is an example of a non-direction-selective Simple cell. In the group of model cells in Fig. 5, cells 1-4 have high DS, cell 6 has some DS while cell 5 has little or no DS. The DS cells in the model have much larger Pref than Opp responses across TF and SF, resembling the cells 1, 2 in the data. Also, they display a great deal of diversity in their tuning curves, as in the data.

**Figure 5.**
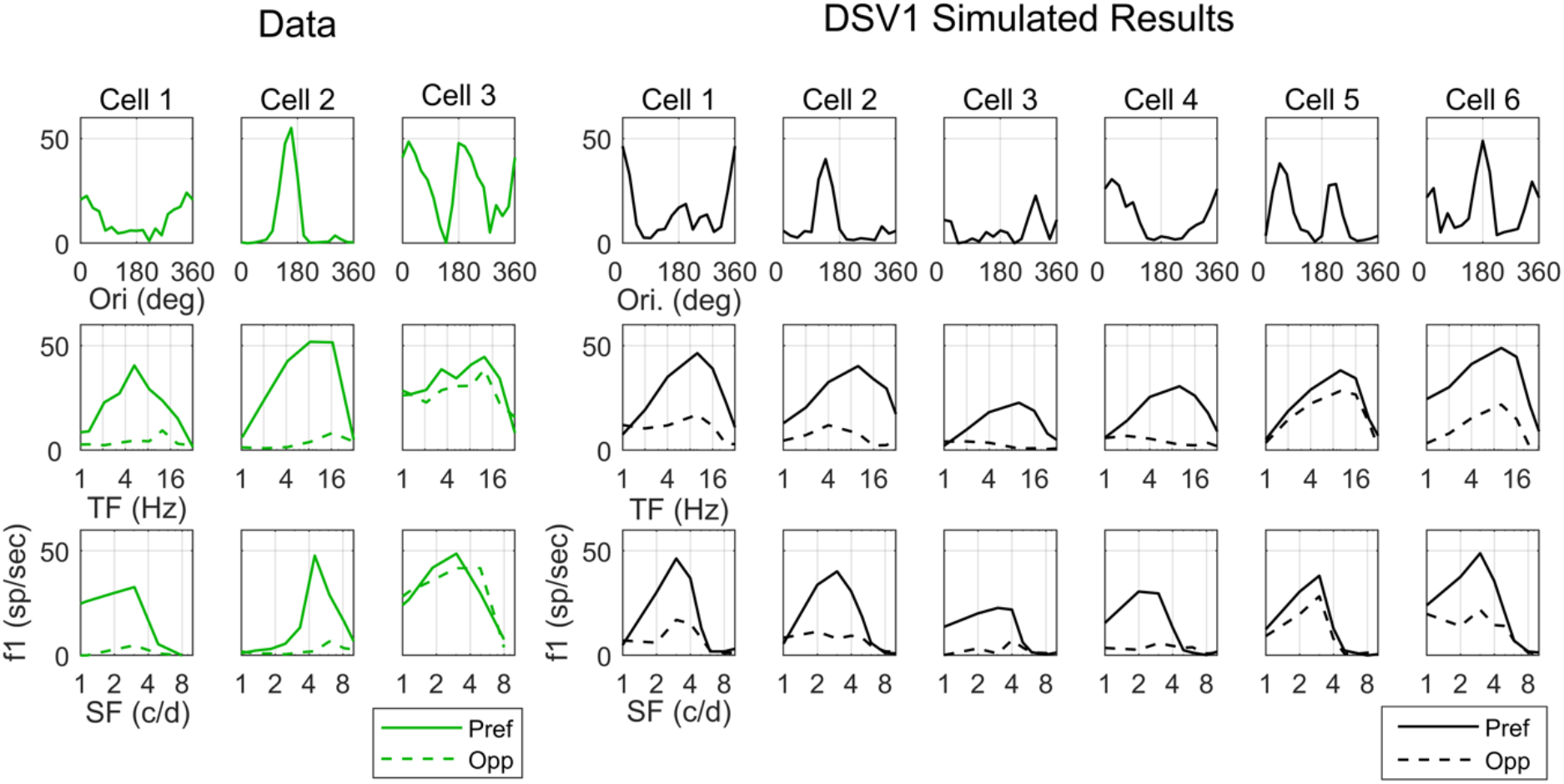
DS properties of single cells from Experimental Data and Model output. Tuning curves of three recorded V1 simple cells (left) and six model simple cells (right) are shown. Top: Orientation tuning curves at optimal TF, SF. Middle: Reponses in Pref and Opp directions vs TF. Bottom: Reponses in Pref and Opp directions vs SF.

### Population data on DS

It is important to compare not only single cell examples but also population data on DS in layer 4Cα with the population data in the DSV1 model.

The first feature of the DSV1 population data is the higher incidence of DS in Simple cells than in Complex cells. For direct comparison with data, Pref and Opp for Simple cells are measured in f1, while f0 is used for complex cells. The fraction of model Simple cells that have Pref/Opp>3 (Figure 6A) is very similar to what is observed in the 4Cα data (Fig. 1C), and the fraction of Complex cells that have Pref/Opp >3 is much smaller, in DSV1 (Fig. 6A) as in the data (Fig. 1C). The higher incidence of DS in Simple cells than in Complex cells (Fig. 6A) is a direct consequence of the structure of DSV1 as explained in connection with Figure 7 below.

**Figure 6.**
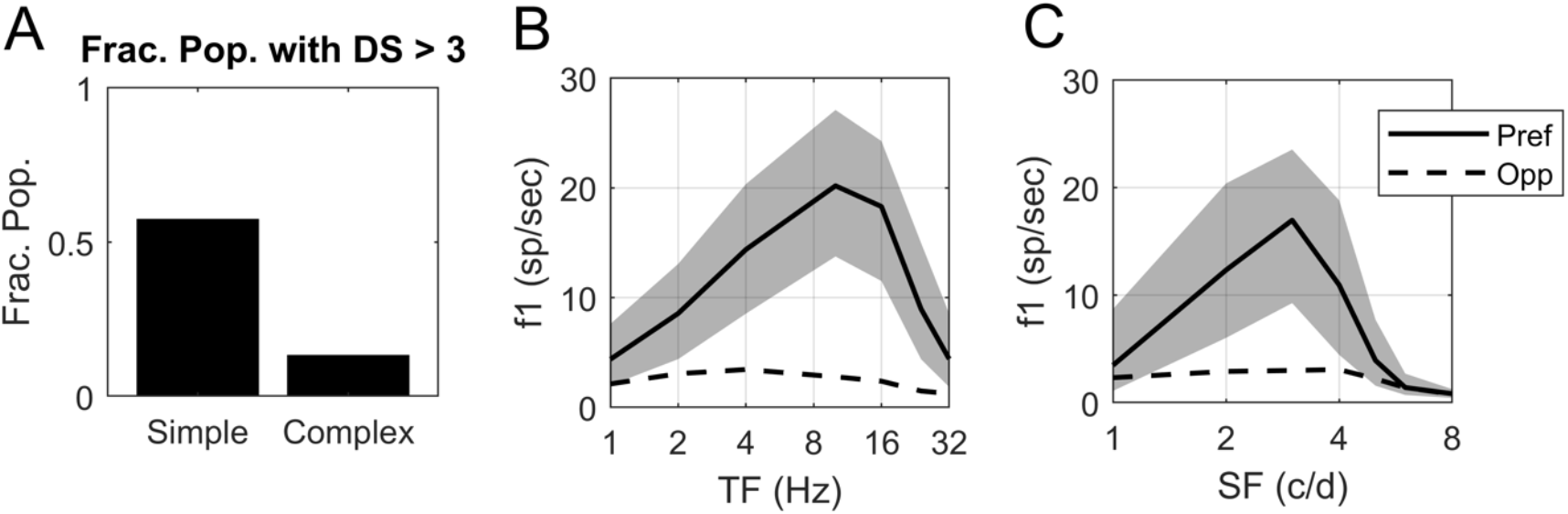
Model data on DS and firing rate as a function of TF and SF. **A**. Fraction of model Simple and Complex cells with Pref/Opp > 3 (compare to Fig. 1C experimental data); the fraction is computed on the population of N=2031 Simple cells and 914 Complex cells in the central hypercolumn of the DSV1 model (Fig. 2A). **B**. Pref and Opp firing rates as a function of TF for those model Simple cells with Pref/Opp>3; N=1164. Thick solid curve shows median Pref firing rate, with quartiles shown by the shaded areas. Dashed curve shows median Opp firing rates. **C**. Pref and Opp firing rates as a function of SF for the same population of 1164 model Simple cells as in **B**. with Pref/Opp>3.

**Figure 7.**
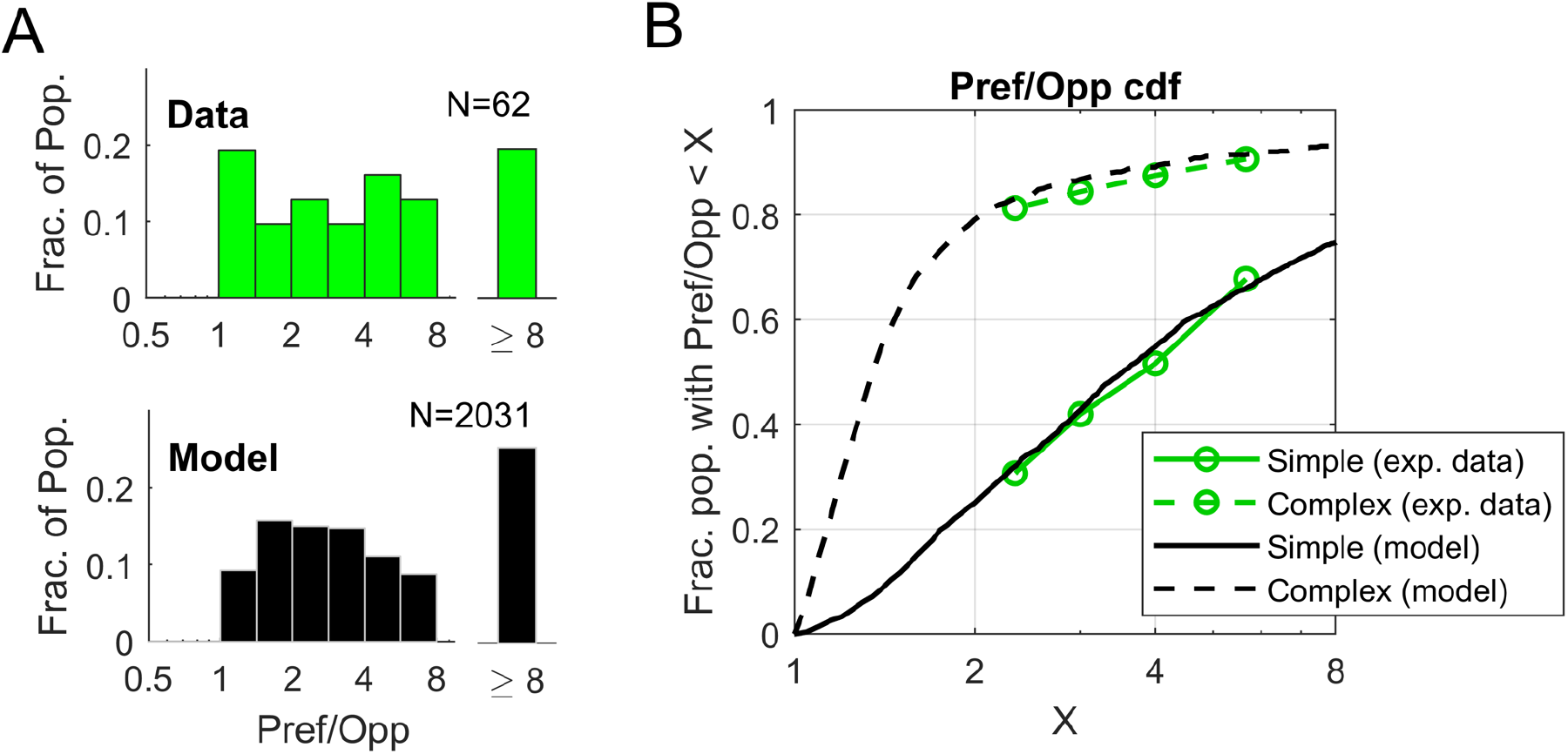
Population statistics of DS in model V1 neurons and comparison with V1 data. **A**. Distribution of DS in model and data for Simple cells. The plots show the distributions of Pref/Opp in model and data for Simple cells for the <Ori, SF, TF> combination that produced maximum f1 response. The data histogram is plotted in green; the model histogram is plotted in black. **B**. Cumulative distributions of Pref/Opp shown for Simple and Complex populations for data (green) and model (black).

A second feature is the comparison of the population average TF and SF tuning in Pref and Opp directions between model and data. As in the data analysis of Fig. 1, “Pref” for a Simple cell is defined as the direction that evokes the cell’s maximal f1 response over all stimulus conditions (orientation, direction, SF and TF); and “Opp” refers to the response in the direction opposite to Pref. The results are that, for responses in the Pref direction, median f1 for model Simple cells (Fig. 6B, C) peaks at TF = 10 Hz and 3 c/deg and these values are similar to those found in the data (Fig. 1D, E, respectively) [cf. Hawken et al 1988; Hawken et al 1996]. As in the data in Fig. 1D, E, the Pref and Opp directions in the model neurons remain the same across TF and SF (Fig. 6B, C). Median responses in the Opp direction are much smaller than Pref responses for model Simple cells (Fig. 6B, C) across TF and SF as in the 4Cα data (Fig. 1D, E). One implication of Fig. 6B, C is that DS in the DSV1 model has the same broad-band character as in the cortical data.

Population distributions of DS across the layer 4Cα population enable a direct comparison of DS in data and model (Figure 7). Figure 7A shows distributions of Pref/Opp in layer 4Cα Simple cells in real cortex (green, upper panel) and in model Simple cells (black, lower panel). As in Figure 1, for each cell in the population, Pref/Opp was computed for the stimulus orientation that produced the largest Pref response. The two distributions in Fig. 7A have very similar shapes. For instance, they have similar medians (∼3.5) and similar fractions of cells with very high DS, i.e. those cells with Pref/Opp>8 (Fig. 7A).

The cumulative distribution functions (CDFs) of DS of Simple and Complex cells in DSV1 simulate those of the data very closely (Fig. 7B). Plotted on the vertical axis of Fig. 7B is the fraction of neurons with Pref/Opp < x, where x is the value of the Pref/Opp ratio on the horizontal axis. The plots indicate that a large fraction of Simple cells in layer 4Cα are directionally selective while Complex cells usually have a small Pref/Opp ratio [Hawken et al 1988; Fig. 6]. The same is true in the DSV1 model. A bootstrapping method was used to test whether or not the CDF of the data was significantly different from that of the model, for both Simple and Complex cell populations, by computing the summed squared deviation between model and data CDFs (see **Materials and Methods** for details). The probability that the Simple cell data were the same as that of the model is p>0.2, while for Complex cells p>0.7. The CDFs of data and model are not distinguishable statistically.

The absence of DS in Complex cells in the DSV1 model (Fig. 6A, Fig. 7B), which is in good agreement with experimental data (Fig. 1C, Fig. 6A), merits comment. As in previous models [Chariker 2016, 2020] the Complex cells in DSV1 typically receive little feedforward input (from 1-2 LGN cells rather than 4-5 for Simple cells), a fact consistent with data [Alonso et al 2001; Hirsch et al 2003]. Since DS in LGN input initiates DS in DSV1 cortical cells, it makes sense that Complex cells without that input should not be direction-selective.

We finish this section with the following two remarks: The first has to do with our use of f1 (as opposed to f0) in measurements of Pref and Opp for Simple cells, which we have done throughout this paper. We have found it natural to use f1 in Layer 4Cα, the input layer to V1, in order to compare V1 Simple cells with their feedforward input. The feedforward input from LGN has DS only in f1: since individual LGN responses are not direction-specific, summed LGN spikes/sec (=f0) are the same in both directions. However, it is different in cortical cell responses; both f0 and f1 responses of V1 Simple cells are direction-selective. In Simple cells in Layer 4Cα, the Pref/Opp ratios computed using f1 and f0 are very similar, both in data and in DSV1. Simple cell CDF’s for model and data, analogous to Fig 7B but using f0 instead of f1, are shown in **Supplementary Information** (S4). CDFs with f1 and f0 are compared there also. They are very similar to each other. That cortical cells also have DS when measured in f0 could be significant functionally, as firing rates may play a role in communication between cortical layers or regions.

Our second remark is on methodology. DSV1 not only has DS cells but it emulates quantitatively the population distribution of DS derived from experiments (Figs. 6 and 7). No other model has achieved this kind of agreement with cortical data on DS at the population level. We want to emphasize that the agreement between model and data was not achieved through curve fitting. DSV1 is a mechanistic model; its network architecture is constrained by known micro-anatomy of V1, its neuronal interactions are designed to simulate those in real cortex, and like its predecessors [Chariker et al 2016, 2018, 2020], its parameters are chosen so that the model –- with a single set of parameters --- replicates simultaneously many V1 phenomena including orientation selectivity, SF and TF tuning, contrast response, and gamma-band oscillations.

### Cortical contribution to DS in the DSV1 model

The mechanisms in [Chariker et al 2021] guarantee that feedforward LGN inputs will have DS, but DS in a V1 cell’s LGN input does not necessarily imply DS in the cell’s response. LGN provides only a small fraction of the mean synaptic current into cortical cells; the bulk of it comes from intra-cortical synaptic transmission (Fig. 4) [cf. Douglas and Martin 2004], and intra-cortical activity could, in principle, modify DS in a cell’s feedforward input. If intra-cortical currents modify a V1 cell’s feedforward-DS, do they enhance or diminish it, and what are the mechanisms? Below we answer these questions for the model. Given the quantitative agreement between DSV1 outputs and data, the findings are testable predictions for the real cortex.

### Comparison of feedforward and cortical DS

The results are summarized in Figure 8

**Figure 8.**
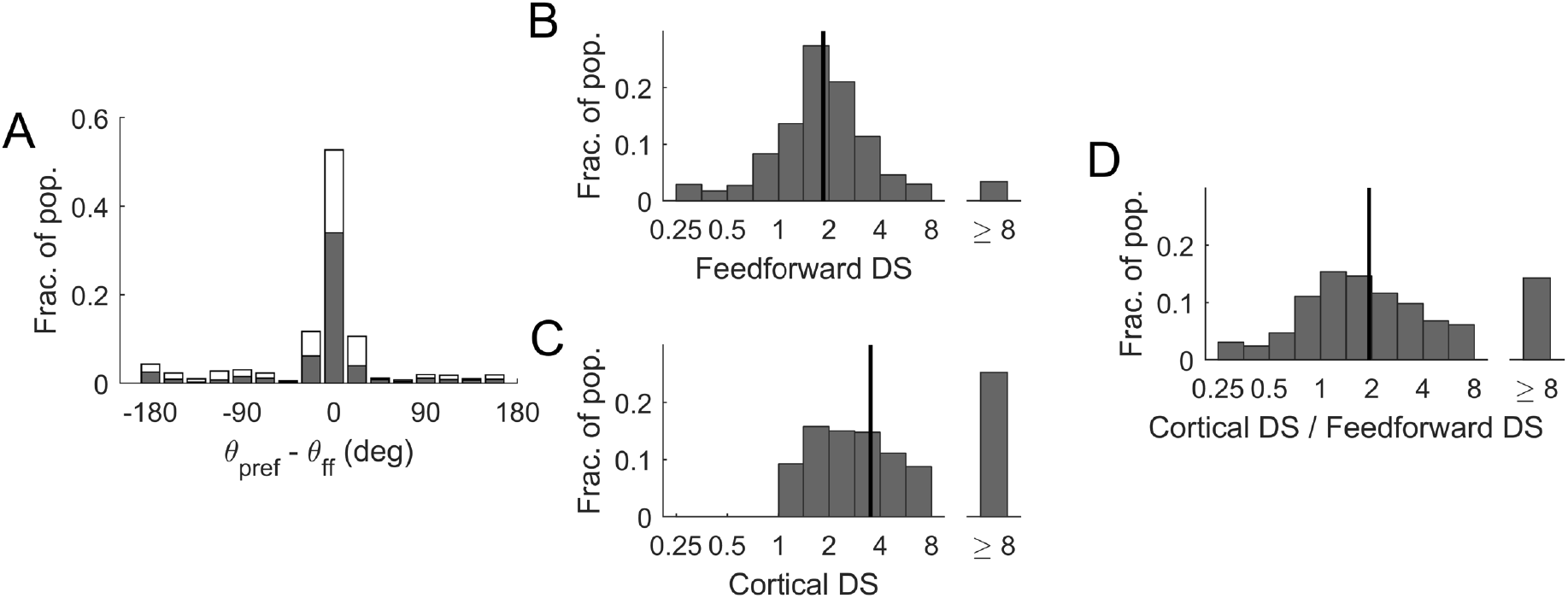
Comparison of DS in feedforward input current and in V1 firing rate for model Simple cells. **A**. Deviations of cortical directional preference from those in feedforward LGN inputs. The statistics shown are for high DS (Pref/Opp>3) Simple cells. Cells with two ON/OFF subregions in their receptive fields (N= 659) are drawn with filled-in bars; cells with three subregions (N=505) are in the unfilled bars. All cells were high DS cells in the central HC of the DSV1 model. The x-axis measures deviation of the cell’s preferred direction from that of its LGN feedforward inputs. **B**. Distribution of DS in the f1 response of the feedforward input current (FF-DS) to Simple cells **C**. Distribution of DS in the f1 response in V1 firing rate in Simple cells **D**. The distribution of the ratio of Cortex-DS/FF-DS cell-by-cell. In B, C, D, the grating that optimizes f1 of the output spike train of the V1 cell is used as Pref in both FF-DS and Cortex-DS. Data are for all Simple cells in the central HC. Black vertical lines are medians; N=2031.

#### Preferred directions

The theory in [Chariker et al 2021] guarantees that for a Simple V1 cell with two receptive field subregions, the preferred direction of its feedforward LGN inputs is from the OFF side to the ON side of the cell’s receptive field. An immediate question is: Is this directional preference passed to the recipient cortical cell? The answer is Yes, as shown in Figure 8A. A majority (about two thirds) of the high DS V1 cells with two subregions followed closely the preferred direction of their feedforward inputs, but about 15% preferred the opposite (Fig. 8A, filled bars). Results for model Simple cells that have three receptive field sub-regions are similar (Fig. 8A, unfilled bars).

#### Magnitudes of DS

DS in the output of a V1 cell, referred to here as *Cortical-DS*, is, as before, defined using as Pref the direction dictated by the stimulus that elicits maximum f1 response in the cell’s firing rate. We will use the same stimulus to define DS in feedforward LGN inputs (abbreviated as *FF-DS*). Explicitly, FF-DS is computed by dividing f1 of the current resulting from the summed spiking LGN input in the Pref direction by f1 of the current from the spiking LGN input in the Opp direction, where the Pref and Opp directions are the same as in the computation of Cortical-DS.

#### Cortex *enhances*

DS above what is present in the feedforward input: the median of Cortical-DS is nearly twice that of FF-DS (Figure 8B, C). This can be seen in the population distributions of FF-DS and Cortical-DS (Fig. 8B, C). Data points are taken from all the Simple cells in the central HC of the DSV1 model. In Fig. 8C, Cortical-DS is ≥1 by definition, but not so in Fig. 8B: FF-DS is < 1 if LGN currents are modulated more in the direction opposite to the preferred direction of the postsynaptic V1 cell; this occurs for a small fraction of the cells, as reported in Fig. 8A.

To give a sense of how FF-DS and Cortical-DS compare on a cell-by-cell basis in the model, Fig. 8D plots a histogram of the ratio Cortical-DS/FF-DS. The histogram shows that while the median of Cortical-DS/FF-DS is around 2, the ratio has a wide spread from well below 1 to more than 8. In other words, while on average the cortical network doubles the Pref/Opp ratio, there is great diversity in the relations between the DS of cortical neurons and of their feedforward inputs.

### Dynamic interaction of LGN and cortical currents

We present below a general theory of current interaction leading to a formula for cortical-DS that can help us understand how cortical interactions affect DS.

#### Synaptic conductances in Pref vs in Opp

In DSV1 there is no directional preference in the synaptic input from other cortical cells. This can be seen by examining the spike input to a model Simple cell from all its E- and I-inputs from other cortical cells in L4 (Fig. 9). The summed rates shown in Fig. 9 are drawn from the population of High DS Simple cells in the central Hypercolumn of the DSV1 model. Observe that both summed E-spike rate (red) and summed I-spike rate (blue) have large DC levels that are approximately the same for Pref and Opp responses, and the same is true for summed spike rates from L6 (not shown). In other words, synaptic input from other cortical cells is not direction-selective. In particular, the summed inhibitory synaptic conductance of high DS cells in DSV1 is not higher in Opp as proposed in some models [Suarez et al 1995; Maex and Orban 1998; Freeman 2021]. For a majority of the cells, there is also no significant dependence of synaptic conductance on LGN phase.

**Figure 9.**
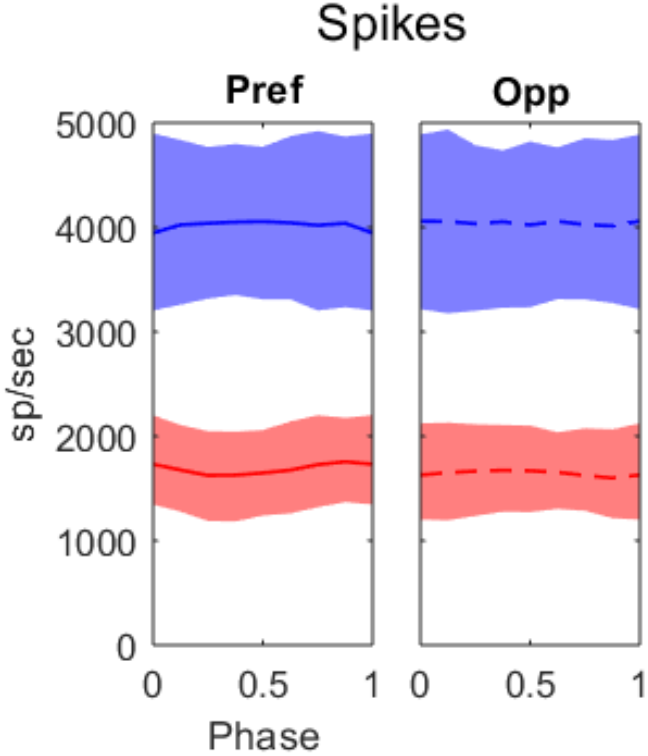
Summed spike rates received by L4 E cells from other L4 cells, in Pref and in Opp directions, plotted against LGN phase (as fractions of a cycle). The E-spikes received are plotted in red, I-spikes in blue. Medians are plotted as the solid (Pref) and dashed (Opp) curves; 25-75% quartiles are indicated by the shading. Data are from high DS cells in the central HC of DSV1; N=1164. The stimuli were drifting gratings with SF=3 c/deg and TF=10 Hz, near the peaks of the population tuning curves (like Fig. 6 B, C). Here and in Fig. 10, LGN phase is depicted as fraction of the stimulus cycle and is in the range [0,1] with 0 corresponding to peak spiking in the LGN input.

#### Phase relations between LGN and cortical currents

While synaptic conductance is flat with phase, the synaptic currents are modulated. The reason is that membrane potential modulation causes modulation of the driving forces of the synaptic currents. For a Simple cell, the membrane potential is modulated at the stimulus frequency f1; during the phase when LGN input is received, membrane potential is more depolarized than during the other half of the stimulus cycle (Figure 10A). In the notation of the LIF equation (see **Materials and Methods**), a more positive value of V at LGN peak phase decreases the driving force for excitation E, (V -V_E_), and increases the driving force for inhibition I, (V – V_I_), where V_E_ and V_I_ are E and I reversal potentials. Because the E and I conductances caused by cortical synaptic connections to a target cell are approximately constant with phase (Fig. 9), the membrane potential modulation causes a significant increase in the magnitude of the I-current into a Simple cell when the LGN input is at its peak and a slightly lowered E-current.

**Figure 10.**
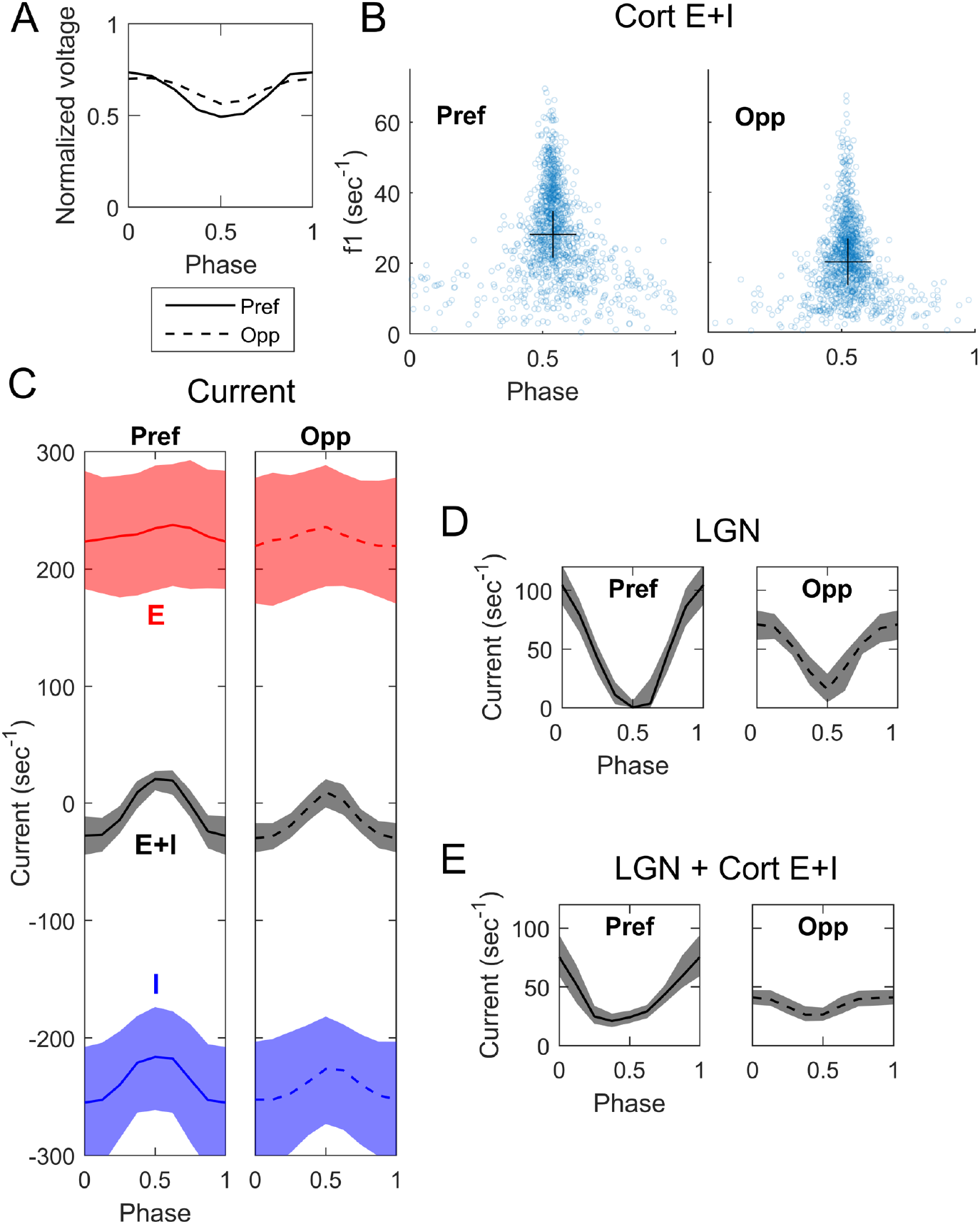
Membrane potential and cortical currents as function of LGN phase in the DSV1 model. Same cell population, stimuli and notation as in Fig 9. **A**. Median modulation of membrane potential in Pref (solid) and Opp (dashed) directions. **B**. The amplitude of the f1 component of net cortical synaptic current (E+I) plotted vs the phase (fraction of a cycle) of the E+I current with respect to the phase of LGN input. Data plotted for the Pref and Opp directions, for each cell in the population. In both directions, the cortical currents are predominantly out of phase (phase ∼ 1/2 cycle) with LGN input. Crosses show medians for both x and y-axes. **C**. E and I synaptic currents vs phase with respect to LGN, in both Pref and Opp directions. E plotted in red, I in blue; medians and quartiles plotted for each. Grey curves and shaded regions are medians and quartiles of the net cortical current, E+I. **D**. LGN excitatory current vs phase with respect to its peak, for Pref and Opp directions. Medians and quartiles plotted as in C. **E**. Sum of LGN and cortical currents, plotted vs phase with respect to LGN.

Through membrane potential modulation, the net cortical synaptic currents for most cells have a tendency to be *antiphase* with respect to the LGN current, and this is true in Pref as in Opp. Figure 10B shows the results for a large population of Simple cells in DSV1. Each point in Fig. 10B is for one cell in the model: f1 amplitude is plotted on the vertical axis and phase is on the horizontal. Phase is plotted as fraction of stimulus period from 0 to 1, 0 being the phase where LGN peaks. The clusters of points near 0.5 phase in both Pref and Opp in Fig. 10B confirm the theoretical expectation that the net (E+I) cortical synaptic currents should be antiphase with LGN input.

#### A formula for cortical-DS

Let {LGN}, {cortex} and {LGN+cortex} denote respectively the f1 of LGN, net cortical, and LGN + cortical synaptic currents into a model cell. Then if net cortical current is antiphase to LGN, we have

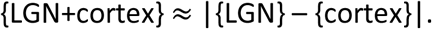

To understand how cortex changes FF-DS, one must examine how the situation differs in the preferred and opposite directions. That is, one must compare

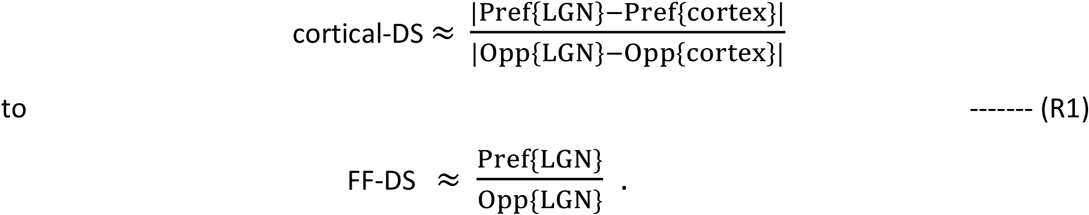

Whether or not Cortical-DS is larger or smaller than FF-DS depends on the relations among the four quantities in eqn. (R1) which we examine next, in Fig. 10 C, D, E.

Consider cortical E and I-currents graphed in Fig. 10C. The black curves with grey shading are the net cortical currents (= E+I), which are anti-phase with LGN, consistent with Fig. 10B. Notice also in Fig. 10C that while E and I-currents each have a large variance, the variance of net cortical current (E+I) is much smaller as a consequence of the very tight balance between cortical E and I-currents for each cell as illustrated earlier (Fig. 4). In Fig. 10D, the excitatory summed LGN currents are graphed vs phase in the cycle, for both Pref and Opp directions, and we can see that they peak at 0 phase (the opposite of the net cortical currents in Fig. 10C). The sum of LGN and cortical currents is graphed in Fig. 10E. One can see in Fig. 10E that the effect of cortical currents is to reduce the synaptic excitation from LGN, for both Pref and Opp directions. In DSV1, cortex reduces Opp current relatively more, and this causes cortical-DS > FF-DS for the population of High DS Simple cells (Fig. 10E vs 10D).

### A detailed study of two subpopulations

To learn more about how the model enhances cortical DS, we study two sub-populations of L4 Simple cells: 1) a sub-population that has DS similar to that of its feedforward input from LGN, and 2) a sub-population that has cortical DS that is magnified compared to its feedforward input. Both populations consist of about 170 Simple cells, and both have 1.4 < FF-DS < 2. Interaction with cortex produces different results, however: cortical-DS < 2.5 for population 1, and cortical-DS > 4 for population 2.

We investigated how the model could generate such different outcomes in these two subpopulations, and observed the following difference in their phase distributions. Fig. 11A and E are scatterplots of phase of net cortical currents in Pref vs Opp directions, using the same meaning of phase as in Fig. 10B. The cells in population 2 almost all have cortical currents with a peak at 0.5 phase (Fig. 11 E) but many cells in population 1 have current peaks significantly different from 0.5, especially in Opp (Fig. 11A).

**Figure 11.**
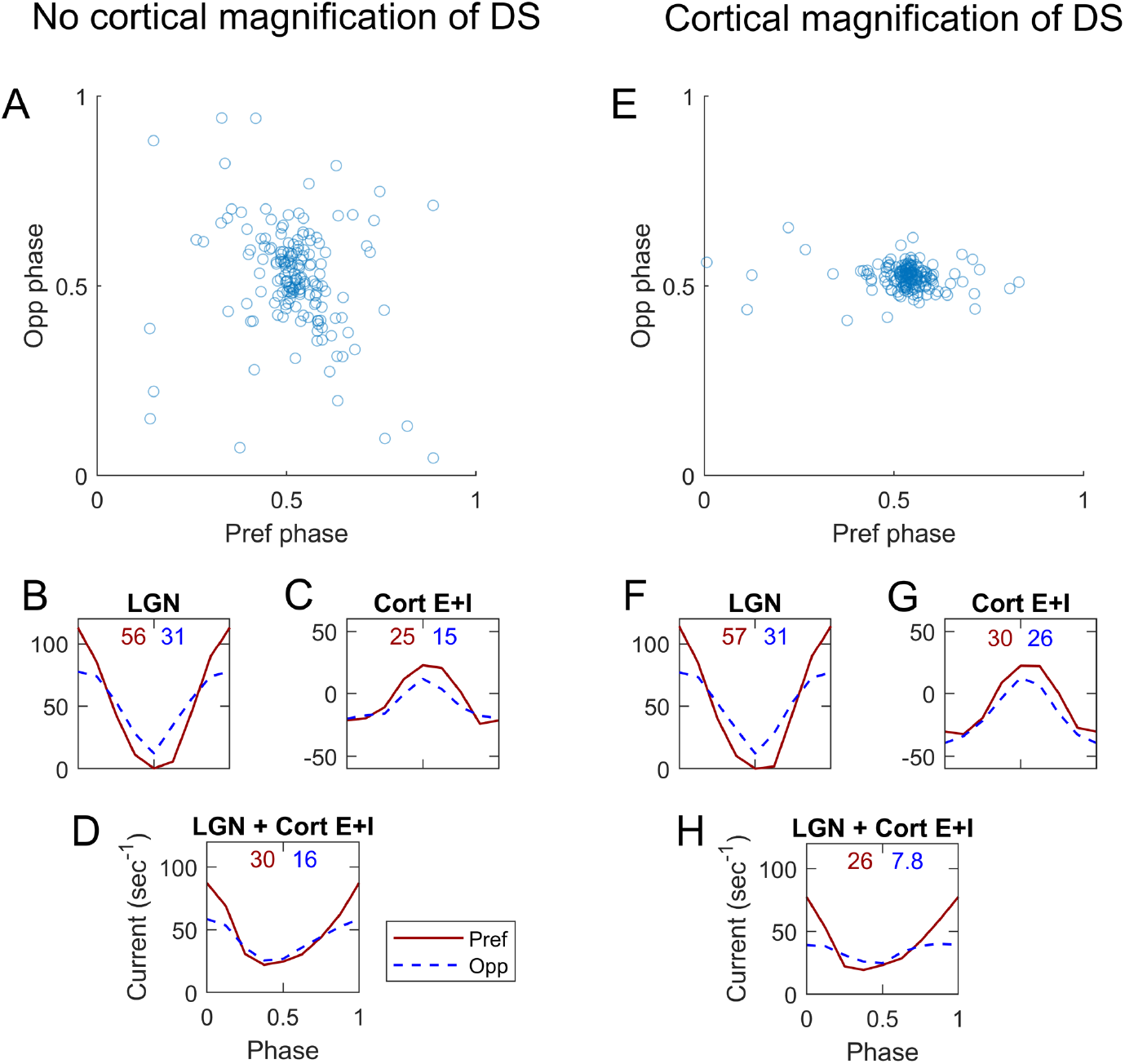
Dynamic interactions between cortical and feedforward synaptic currents in two groups of model L4 Simple DS cells. **A-D** from a group of 167 L4 cells that inherit all their DS and do not enhance it. **E-H** from a different group of 176 L4 cells, cells that enhance DS well above that in the LGN input. Both groups were selected to receive the same Pref/Opp ratio of ∼1.8 in their LGN inputs. **A**. Phase of membrane current vs LGN phase in response to a drifting grating <SF=3 c/deg, TF=10Hz>. Opp phase is plotted vertically, and Pref phase horizontally. Note the wide spread of the phases especially in Opp (Std. dev.=0.14 in the Opp direction). **B**. Medians of the cycle averages of LGN synaptic current; Pref (solid), Opp (dashed). F1 (in normalized units of sec^-1^) for the median are written above the plotted curves. **C**. Corresponding median synaptic currents (E+I) for L4 cells in Pref and Opp directions. **D**. Medians of the sums of LGN and cortical synaptic currents in Pref and Opp directions. The Pref/Opp ratio =1.9 is roughly the same as for the LGN input in **B**. **E-H** are analogous to **A-D** for the second group of cells. Note that in **E**, the Std. Dev of the phase in the Opp direction (=0.03) is much less than in **A**, and the Pref/Opp ratio (=3.3) of the Summed LGN + Cort E+I current is higher in **H** than in **D**.

As to how this is related to cortical magnification of DS, the answer lies in our previous analysis of the dynamics of cortical interactions. Fig. 10A shows that modulations of the membrane potential *V* of a Simple cell follow the cell’s feedforward LGN inputs, and that *V* has a minimum at midphase (i.e. at 0.5 cycle) with respect to LGN. When incoming spikes from cortex have little or no phase preference, cortical current peaks at ∼0.5 phase as a consequence of the minimum of *V*. A cortical current peak at ∼0.5 phase is the case for a majority of High DS cells in DSV1 (Fig. 10B), and also for almost all cells in population 2 (Fig.11E). But when recurrent spike rates vary with phase, their synaptic inputs could interfere with the feedforward modulation of *V* and, in this way, could compromise the f1 of cortical current as well as cause its preferred phase to be displaced.

The difference in recurrent dynamics appears to be what is happening in the two populations in the Opp direction: phases in population 1 are more diffused than in population 2 (Fig. 11A, E), and the f1 of (cortical E+I) in population 1 is also smaller than that in population 2 (Fig. 11C, G). Differences of f1 in Pref are less significant. Thus, in population 1, interaction with cortex leads to the subtraction of a smaller number from the denominator in Eqn (R1) compared to population 2. Subtraction of a larger number from the denominator causes the resulting Pref/Opp ratio to be larger, hence cortex magnifies DS more in population 2.

## Discussion

Motion perception is a vital visual capability for primates including humans. We use visual motion perception to track objects, to break camouflage, to identify surfaces and objects, and to navigate [Nakayama 1985; Ullman 1988; Borst and Euler 2011]. Perceiving the direction of motion is one crucial element of motion perception and that’s why visual neurons with DS constitute the neural basis of motion perception [Parker and Newsome 1998]. This is our motivation to achieve a theoretical understanding of the neural mechanisms of DS in the visual cortex.

## The DSV1 model: aims and predictions

### Aims of study and summary of findings

Based on **Results**, we answer the two questions posed at the end of **Introduction**. First, with the construction of DSV1, we demonstrated that the theory of Chariker et al [2021] could be implemented in a biologically-realistic, large-scale network model. The sparseness of LGN posed significant challenges in LGN template design (see **Model Description**) but the templates did produce FF-DS (Fig. 8B). Second, our results dispelled the concern that DS in the feedforward input may not be enough to generate significant DS in the firing rates of V1 neurons: Comparisons of DSV1 output with experimental data (Figures 5-7) show that the ON/OFF hypothesis alone, without the aid of intra-cortical connections designed specifically to promote DS, could initiate the amount and quality of DS seen in V1 data. Though LGN input current is a small fraction of intra-cortical currents in V1 [Douglas and Martin 2004], we found that (i) E and I-currents within cortex are so tightly balanced that LGN inputs, which break the balance, have a large effect (Fig 4), and (ii) dynamic interaction of feedforward and intra-cortical currents leads to DS enhancement (Fig 10).

We do not claim that the mechanisms proposed in this paper are the only mechanisms that contribute to DS. The presence of specific intra-cortical connections would not contradict our theory. What we have shown is that the magnitude and quality of DS seen in data can be produced without such connections.

### Model predictions

One of the purposes of a theory is to stimulate experiments to confirm or refute its predictions. Here are some specific predictions of the DSV1 model:

1. the preferred direction of a DS layer L4 Simple cell with two ON-OFF subfields should be motion from the OFF side to the ON side of the receptive field (Figure 8A);
2. the physical distance *d* between rows of ON and OFF LGN cells in the LGN input to a V1 cell determines the range of SF to which a cortical cell responds with a consistent directional preference (Figures 5, 6);
3. on average across the cortical population, the DS of a L4 cell is much greater than the DS in its feedforward input but there is a wide diversity in the degree of enhancement of DS in the cortex (Figures 8 and 10);
4. net cortical inhibition, in antiphase with feedforward excitation from LGN, enhances cortical DS (Figures 10, 11).

### Context: other models

The significance of the DSV1 model can be understood in the context of many earlier models of Direction Selectivity (DS). The earlier models fall into two main groups: 1) feedforward models, and 2) intra-cortical models.

#### Feedforward Models of DS

Many investigators have proposed, as we do, that spatio-temporal inseparability (STI) in the feedforward input to the cortex initiates cortical DS [McLean and Palmer 1989; Priebe and Ferster 2005; Saul et al 2005; Lien and Scanziani 2018; Billeh et al 2020]. But the tough theoretical question is, what are the neural mechanisms responsible for STI?

One early proposal was that there are different types of LGN cells, so-called lagged and non-lagged cells, with very different temporal kernels [Saul and Humphrey 1992]. The lagged-cell mechanism has been invoked to explain DS data in cat [Saul and Humphrey 1992] and ferret [Moore et al 2005] V1. While the motion-energy model [Adelson and Bergen 1985] was not concerned with biological implementation, later work on biological mechanisms of the motion-energy model proposed that the lagged-cells could be the source of STI [Emerson et al. 1992]. However, lagged cells have not been found in Macaque LGN [Wiesel and Hubel, 1966; Kaplan and Shapley, 1982; Hicks et al 1983; Derrington and Lennie, 1984] and therefore the source of DS in Macaque V1 must come from other mechanisms [Saul et al 2005]. Furthermore, the lagged cell mechanism would have trouble explaining the broad-band nature of DS in Macaque especially the high value of the Pref/Opp ratio at high TF (Fig. 1D).

A number of studies have suggested that STI leading to DS comes from the differential timing of Transient and Sustained input to DS cortical cells [Marr and Ullman 1981]. Some experimentalists suggest [McLean and Palmer 1989; Priebe and Ferster 2005}, and some theories propose, Sustained and Transient LGN-->cortex connections [Lien and Scanziani 2018; Billeh et al 2020] as the source of cortical DS. One problem for Sustained-Transient models in Macaque is that there is no evidence for a Sustained type of a Magnocellular input to Macaque V1 layer 4Cα [Derrington and Lennie, 1984; Reid and Shapley 2002; Saul et al 2005]. A second problem that has not been answered by Sustained-Transient models: how can a cortical cell select Sustained LGN input for one receptive field sub-region and Transient LGN input for a spatially-distinct sub-region? This is the wiring problem that is solved by the OFF-ON hypothesis [Chariker et al 2021].

#### Intra-cortical models

Others proposed that the feedforward input to the cortex is not DS but rather that there was, initially, the generation of non-DS Transient and Sustained cortical RF’s in cortical cells [DeValois et al, 2000; Baker and Bair 2012]. When these cortical cells’ activity was combined intra-cortically it would give rise to DS [DeValois et al, 2000; Baker and Bair 2012]. There are two distinct problems with the specific models proposed. DeValois et al [2000] proposed that the Sustained branch of their model was comprised of cortical Simple cells that received Parvocellular feedforward input, while the Transient branch was cortical cells excited by Magnocellular LGN cells. This is not plausible for cells in layer 4Cα that receive predominantly Magnocellular LGN input [Lund 1988; Chatterjee and Callaway 2003; Angelucci and Sainsbury 2006]. Furthermore, DS is present at low contrast [Saul et al, 2005], and only Magnocellular cells provide useable responses at low contrast [Kaplan and Shapley 1982; Hicks et al 1983; Derrington and Lennie 1984]. The model of Baker and Bair [2012] resembles that of DeValois et al [2000] without being species-specific. Neither model answers the question specific to layer 4Cα, namely how is DS produced in such abundance (Fig. 1C, Fig.7) at such an early stage in cortical signal processing?

Another very different kind of intra-cortical model of DS is that the excitatory drive of a DS cell is not DS but rather that inhibitory cortico-cortical input selectively suppresses responses in the non-preferred direction [Suarez et al 1995; Maex and Orban 1998; Freeman 2021]. This kind of theory is not consistent with experimental data that indicate that intra-cortical inhibitory conductance is somewhat higher amplitude in the Pref direction [Priebe and Ferster 2005].

### Modeling approach -- compared with Machine Learning

It is instructive to compare our modeling approach to machine learning (ML), an increasingly important investigative tool in neuroscience. ML involves training large neural networks such as Convolutional Neural Networks, or CNNs, to perform specific tasks [e.g. Bashivan et al, 2019; Zhang et al, 2019]. The strength of ML lies in the fact that it can deal with high-dimensional complexity and requires no *a priori* knowledge of anatomy or of how the brain performs the tasks. The price one pays for this ``black-box” type approach that focuses on input-output relations is that they are not designed to inform about mechanisms, and extracting such information can be extremely challenging. We have, in this paper, employed a bottom-up approach. In DSV1, which employs 36,000 model neurons, there is a direct correspondence between anatomical and model structures. Model components and functions were benchmarked to provide a comparison with their counterparts in real cortex. Consequently, one can expect that the dynamics of the DSV1 model are a good reflection of what goes on in real cortex; they have the potential to explain the neuronal mechanisms.

We note finally that the usefulness of a biologically realistic model goes beyond the model itself: it provides a well-constrained feedforward component to the next stages of processing in cortex. As layer 4Cα provides the dominant input to layer 4B in V1, DSV1 will provide an essential contribution to the modeling of layer 4B, and potentially even to the modeling of extra-striate cortical areas that receive input from layer 4B such as the thick stripes of V2 [Sincich and Horton, 2005; Federer et al, 2009] and MT [Sincich and Horton, 2005].

## Acknowledgements

This research was supported by NIH Grant R01 EY001472 (R.S.), NIH Grants R01 EY008300 and NIH R01 EY 15549 (M.H.), NSF Grant 1734854 (R.S. and L.-S.Y.), NIH Core Grant P30 EY13079, and NIH Training Grant T32 EY7136. We thank the graduate students and postdoctoral fellows who participated in the experiments—Dario Ringach, Michael Sceniak, Elizabeth Johnson, Siddhartha Joshi, J. A. Henrie, Patrick Williams, Dajun Xing, Christopher Henry, and Anita Disney — and Madhura Joglekar for early numerical explorations.

