## Supplementary material for "A computational model of direction selectivity in Macaque V1 cortex based on dynamic differences between ON and OFF pathways": Supplmentary Information

### Supplemental Information

#### S1 LGN sustained component and saturation of firing rate

**LGN sustained component.** The step response of an ON cell with temporal kernel of type $(a, b)$, defined in Results, will have a sustained component when $a \neq b$, corresponding to the nonzero total area of the kernel (a $(1, 1)$ kernel has zero total area). In the distribution of temporal kernels given to model ON cells, shown below, one can see that all kernels have nonzero total area.

| Kernel | $(1.7, 0.8)$ | $(1.6, 0.7)$ | $(1.1, 0.5)$ | $(1.0, 0.4)$ |
|---|---|---|---|---|
| Probability | 10% | 30% | 30% | 30% |

ON cell kernels derived from experiment (Reid and Shapley, 2002) show that ON cells can have kernels with either zero or nonzero total area, and therefore in our model, for 1/2 of all ON cells, we horizontally stretch the area after the zero crossing to make the total area equal to zero. This modification of an $(a, b)$ kernel can be written explicitly

$$K_{a,b}^{\text{mod}}(t) = \begin{cases} aK_{\text{OFF}}(t), & t \leq 35, \\ bK_{\text{OFF}}(b(t-35)/a), & t > 35. \end{cases}.$$

A consequence of this modification for direction selectivity (DS) is that the DS of the feedforward input provided by a pair of ON/OFF LGN cells is reduced at low temporal frequencies (TFs). Figure S1 shows the DS in the summed output of a pair of ON/OFF LGN cells, with a setup as in figure 2C of the main text, in the case of an unmodified (nonzero total area) and modified (zero total area) LGN cell. At low TF, less than $\sim$ 4 Hz, DS of the output can be seen to be significantly reduced.

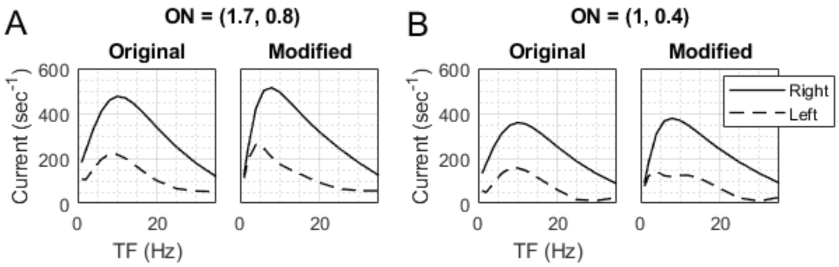


Figure S1: **Effect on DS of removing ON cell sustained component.** The summed response of a pair of ON and OFF LGN cells is shown as a function of temporal frequency, with setup idential to figure 2C of the main text. The ON cell is placed $0.1°$ to the right of the OFF cell, and both left- and right-moving grating response are shown. Grating spatial frequency is 2.5 c/d. The OFF kernel type is (1,1). The ON kernel is type $(1.7, 0.8)$ in (A) and $(1, 0.4)$ in (B). Both the original kernel, with nonzero total area, as well as the modified kernel, with zero total area, are shown. In all cases, ON cells have a 10 ms delay added.

**Firing rate saturation.** LGN cell firing rate is known to saturate under strong drive (Kaplan et al, 1987). To make it harder to spike at high firing rates, we implemented a time-varying spiking threshold which instantaneously jumps up a fixed increment after each spike and relaxes back towards a baseline value inbetween spikes. If $t_j^{(i)}$, $j = 1, 2, \ldots$, are the spike times of cell $i$, then the spiking threshold $V_i^{\mathrm{th}}$ of that cell follows the equation

$$\frac{d}{dt} V_i^{\mathrm{th}} = -\frac{1}{\tau^{\mathrm{th}}} (V_i^{\mathrm{th}} - 1.2) + S^{\mathrm{th}} \sum_{j=1}^{\infty} \delta(t - t_j^{(i)})$$

where $S^{\mathrm{th}} = 0.3$ and the value of the relaxation time $\tau^{\mathrm{th}}$ depends on the number of recent spikes. The relaxation time is longer when the number of recent spikes is larger, further penalizing a high firing cell. If $R_i(t)$ is the number of spikes $t_j^{(i)}$ such that $t - 18 < t_j^{(i)} < t$, then the relaxation time is given by the following table:

| $R_i(t)$ | 0 | 1 | 2 | 3 | $\geq 4$ |
|---|---|---|---|---|---|
| $\tau^{\mathrm{th}}$ | 15 ms | 15 ms | 22.5 ms | 37.5 ms | 50 ms |

Figure S2 shows the effect of the saturation mechanism on LGN firing rate as a function of temporal frequency. It shows that firing rate is attenuated more strongly at the preferred temporal frequencies of the LGN cell.

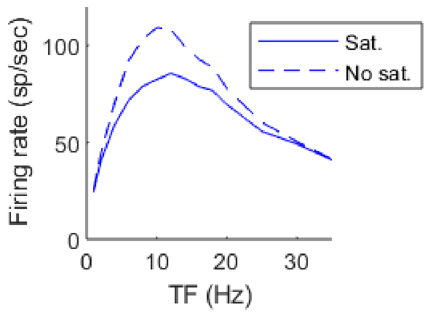


Figure S2: **Effect of saturation mechanism on a (1,1) LGN cell.**

#### S2 LGN templates

The LGN templates with 4-6 inputs are shown in figure S3. Each template has a list of attributes, and it is explained below how each of these attributes is to be distributed over the L4 E-cell population when assigning templates to E-cells:

1. **Number of inputs.** The number of LGN inputs, $n_{\mathrm{LGN}}$, to an E-cell is a random number sampled from the distribution below:

| $n_{\mathrm{LGN}}$ | 1 | 2 | 3 | 4 | 5 | 6 |
|---|---|---|---|---|---|---|
| Prob. | 10.5% | 20% | 2% | 21% | 32.5% | 14% |

2. **Number of ON/OFF stripes.** For E-cells with 4-6 inputs, 2/3 are assigned templates with 2 ON/OFF stripes, and the remaining are assigned 3 stripes.

3. **Template categories.** The templates are divided into two categories: a primary one labeled "Main" used by 80% of E-cells, and a secondary one labeled "Other" used by the remaining 20% (sampled randomly). The "Other" category contains templates exceeding the upper bound on ON/OFF separation $d$ as explained in part I.

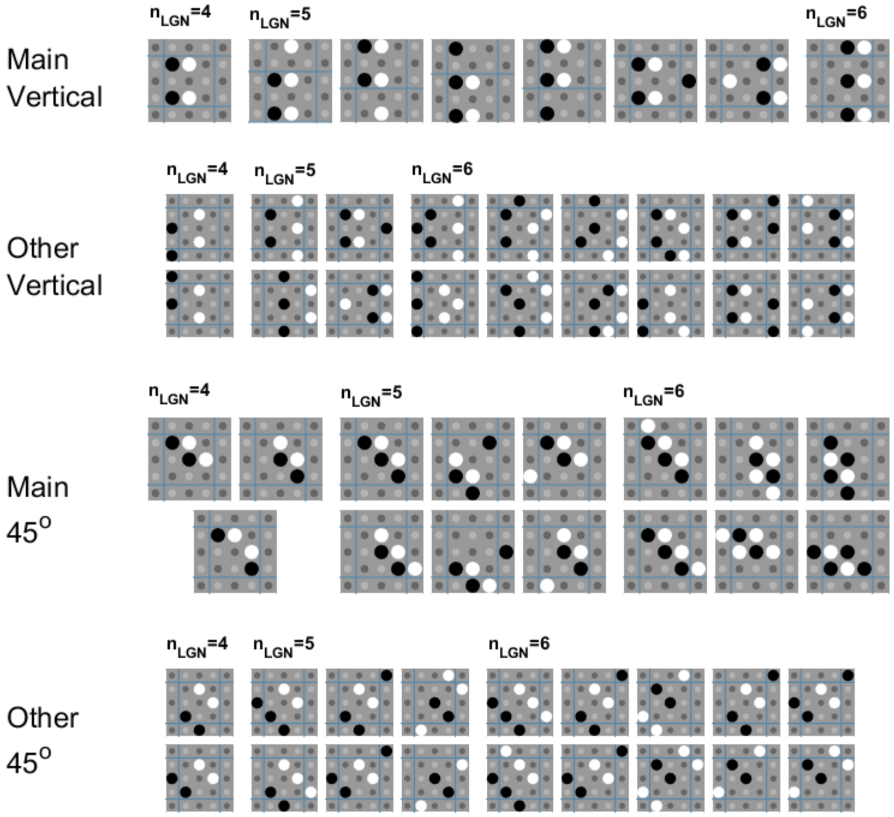


Figure S3: LGN templates. Shown are the LGN templates with 4-6 inputs, vertical and 45$^o$ orientations, sorted into collections labeled "Main" and "Other".

### S3 New Features of DSV1

For a more detailed description of model features which are not specifically related to DS and largely unchanged, see Chariker et al (2020) and its supplemental information.

**Firing rate homeostasis.** We gave simple cells with high firing rate more I-inputs. To determine which cells will receive extra I-inputs, we first measured for each L4 E cell $i$ its firing rate $R_i$ optimized over a range of spatial and temporal frequencies and orientations. The following table gives the exact number of I inputs added as a function of $R_i$:

| $R_i$ | 0-20 | 20-35 | 35-48 | $\geq 48$ |
|---|---|---|---|---|
| Change in # I inputs | -1 | 0 | +3 | +9 |

**Adaptation at low temporal frequencies.** To prevent an excess buildup of complex cell firing rate at low temporal frequencies, we implemented an adaptation mechanism which works by boosting the effectiveness of the I-inputs for any L4 E-cell which spikes enough times in quick succession. A chain of 7 or more spikes with interspike times no longer than $\sim$ 24-27 ms in-between them activates the mechanism, and it becomes deactivated when the cell stops spiking for more than that duration. While adaptation is activated, every I-spike received by the E-cell comes with an additional slow time course

$$G^{\text{adapt}}(t) = S^{EI}\left(\frac{1}{\tau^{\text{decay}}}\exp\left(-\frac{t}{\tau^{\text{rise}}}\right) - \frac{1}{\tau^{\text{rise}}}\exp\left(-\frac{t}{\tau^{\text{rise}}}\right)\right)$$

with time constants $\tau^{\text{rise}} = 5$ ms and $\tau^{\text{decay}} = 100$ ms, and with synaptic strength $S^{EI}$ equal to the usual I-to-E strength. This time course resembles that of $\text{GABA}_\text{B}$, known to be present in V1 (Zilles et al., 2004). If the cell spikes 8 or more times, then the strength of the slow time courses is doubled while adaptation is active.

#### S4 Direction selectivity of mean firing rate

In Results, the direction selectivity of a Simple cell in the model was defined to be the ratio between its f1 responses in the Pref and Opp directions. This was done to compare with experimental data, where the f1 responses were also used to compute DS (see Materials and Methods). Here we compute the direction selectivities of the mean firing rate responses as well.

For each Simple cell, let $\text{Pref}^{\text{f0}}$ and $\text{Opp}^{\text{f0}}$ be the f0 responses of the cell when driven by its Pref and Opp gratings (the same gratings as defined in Results). We then define the DS of its f0 response to be $\text{Pref}^{\text{f0}}/\text{Opp}^{\text{f0}}$. In the left panel of figure S4, the distribution of Simple cell DS is recomputed this way (compare to figure 7B in the main text, where DS in f1 is shown for Simple cells). It shows that there is significant DS in the mean firing rate responses, and that the model and data have similar DS distributions. The right panel of figure S4 shows a comparison in the model between the distributions of DS in f0 and DS in f1.

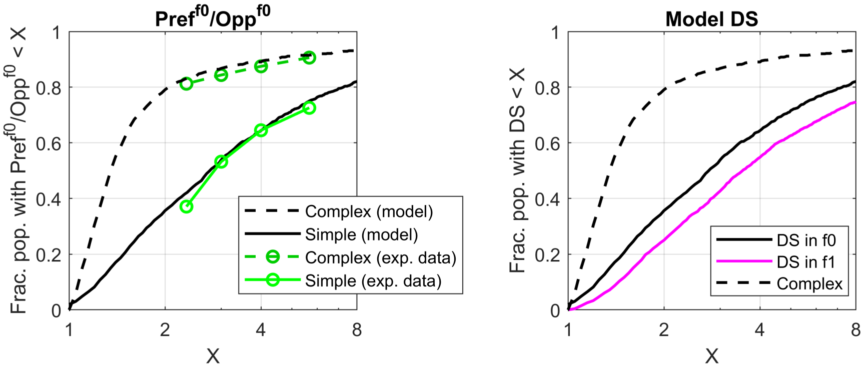


Figure S4: Distribution of DS of mean firing rate in both the model and data. In the left panel, the distribution of DS of the mean firing rate is shown for both Simple and Complex cells, both for model and data. In the right panel, the Simple cell distribution of DS in f1, as it was computed in the main text, is added in purple for comparison.

#### S5 Parameters used

The following table gives values for model parameters. See Chariker et al. (2020) supplemental for more parameters.

| Parameter | Value | Description |
|---|---|---|
| $\tau_{\text{leak}}$ | 0.1 $\text{ms}^{-1}$ | LGN cell membrane leak rate (see $V(t)$ eq. in Methods) |
| $I_B$ | 0.1 $\text{ms}^{-1}$ | LGN background current (see $I(t)$ eq. in Methods) |
| C | 0.109 | Input current prefactor (see $I(t)$ eq. in Methods) |
| $S^{EE}$ | 0.0230 | E→E synaptic strength |
| $S^{EI}$ | 0.0497 | I→E synaptic strength |
| $S^{IE}$ | 0.0059 | E→I synaptic strength |
| $S^{II}$ | 0.0321 | I→I synaptic strength |
| $S^{ELGN}$ | 0.0529 | LGN→E synaptic strength |
| $S^{ILGN}$ | 0.0655 | LGN→I synaptic strength |
| $S^{EL6}$ | 0.0080 | L6→E synaptic strength |
| $S^{IL6}$ | 0.0020 | L6E→I synaptic strength |
